# Structural insights into the formation of repulsive Netrin guidance complexes

**DOI:** 10.1101/2022.09.03.506478

**Authors:** Jessica M. Priest, Ev L. Nichols, Juan L. Mendoza, Kang Shen, Engin Özkan

## Abstract

Netrins can dictate attractive and repulsive responses during axon growth and cell migration, where presence of the receptor UNC-5 on target cells results in Netrin-mediated repulsion. Molecular details of Netrin–UNC-5 interactions and how they signal remain elusive. Here, we show that nematode UNC-5 is a heparin-binding protein, and the UNC-5–heparin affinity can be modulated using directed evolution or via rational design using our novel structure of UNC-5 with a heparin fragment. Furthermore, UNC-5 and nematode UNC-6/Netrin form a large, stable and rigid oligomeric complex in the presence of heparin, which can incorporate the attractive UNC-40/DCC receptor, demonstrating binary and ternary ectodomain complexes at preparative scale. *C. elegans* with a heparin-binding deficient UNC-5 fail to establish proper gonad morphology due to abrogated distal tip cell migration, which relies on repulsive UNC-5 signaling in response to UNC-6. Our findings establish Netrin responses to be mediated through glycosaminoglycan-regulated large macromolecular complexes.

## INTRODUCTION

Synaptic connectivity of the nervous systems in bilaterian animals is established through a conserved set of wiring molecules, including those that guide the growth of axons and determine synaptic partners (Cortés et al., 2022; Kolodkin and Tessier-Lavigne, 2011; Zang et al., 2021). Netrins are conserved, secreted guidance cues that are unique in their ability to exert attractive and repulsive responses on growing axons (Boyer and Gupton, 2018). In addition to their neuronal functions, Netrins and their receptors are known to control cell proliferation, migration, differentiation and survival, and thus they are targets for cancer treatments and insulin resistance (Brisset et al., 2021; Ko et al., 2012; Ramkhelawon et al., 2014).

Netrins, known as UNC-6 in nematodes, act through two classes of receptors: DCC/Neogenin class of receptors (named UNC-40 in nematodes) are required for both attractive and repulsive responses, while UNC-5 receptors are required only for repulsive responses (Hedgecock et al., 1990). For attractive signaling, Netrins can cause oligomerization or clustering of DCC or Neogenin through the interaction of the receptors with multiple sites on Netrin molecules (Finci et al., 2014; Xu et al., 2014). Such oligomerization likely leads to the proximity of intracellular signaling molecules recruited at motifs conserved in DCC/Neogenin cytoplasmic domains (Meijers et al., 2020). It is not clear, however, how Netrin can induce repulsive responses using both DCC and UNC-5-like receptors, as there is only limited biochemical characterization and structural information for the proposed Netrin–UNC-5 complexes.

Another important player in the Netrin guidance system is glycosaminoglycans, which are present on cell surfaces in the form of proteoglycans. Netrins were first purified from tissue with the help of their strong affinity for heparin (Serafini et al., 1994). Their relevance for signaling has also been recognized, as lack of the heparan sulfate-polymerizing enzyme, Ext1, leads to defective Netrin- and DCC-mediated axon guidance in mouse spinal cord commissural neurons (Matsumoto et al., 2007). Mutations in two heparan sulfate proteoglycans (HSPG), syndecan (SDN-1) and glypican (LON-2), show defects in UNC-6/Netrin-mediated repulsive dorsal guidance of motor axons in *C. elegans*. A screen searching for mutations to enhance HSPG-dependent guidance pathway defects identified multiple *unc-5* mutants, hinting at an interaction between UNC-5 and HSPGs (Gysi et al., 2013). Similarly, mutations in the HSPG *unc-52/perlecan* enhance distal tip cell migration defects of *unc-5* hypomorphs (Merz et al., 2003). An interaction between *C. elegans* UNC-40/DCC and the protein backbone of LON-2/glypican, proposed as a modulator of Netrin-mediated axon guidance, has also been reported (Blanchette et al., 2015). Finally, Netrins can act as short-range or immobilized cues (Brankatschk and Dickson, 2006; Moore et al., 2009; Varadarajan et al., 2017), where Netrins would need to be immobilized within the growth substrate (Moore et al., 2012). Interactions with cell surface proteoglycans or the extracellular matrix can immobilize Netrin and allow for signaling via mechanotransduction and haptotactic responses.

Given clear functional relevance and available structural models, Heparan sulfate (HS) binding has been proposed as a mechanism for regulating receptor selection and binding to Netrins (Meijers et al., 2020), although experimental evidence is lacking. It is not clear if the contribution of glycosaminoglycans is essential for signaling, as the major heparin-binding site of Netrins, the C-terminal NTR domain (Kappler et al., 2000) is not required for guidance activity of UNC-6/Netrin (Lim et al., 1999) or binding to either DCC/Neogenin or UNC-5 (Keino-Masu et al., 1996; Leonardo et al., 1997). As Netrin without the NTR domain might still have some heparin-binding activity, it has been difficult to dissect the roles of glycosaminoglycans in Netrin signaling via simple domain deletions and truncations.

Finally, the functional roles of HSPGs in axon guidance is not limited to Netrin signaling. For example, heparan sulfate is known to mediate interactions between other guidance cue-receptor pairs, including Slit and Robo (Hussain et al., 2006), and the LAR class of neuronal receptor tyrosine phosphatases to control neurite outgrowth (Fox and Zinn, 2005; Johnson et al., 2006). Heparan sulfate is also shown to interact with classical guidance cues Semaphorins and Ephrins (Shipp and Hsieh-Wilson, 2007).

While Netrin’s affinity for heparin-like molecules is well established, it is not clear if the receptors interact with glycosaminoglycans directly (Geisbrecht et al., 2003). Here we chose to work with *C. elegans* UNC-6/Netrin and UNC-5, the first molecules of their families to be identified, since they have clear axonal guidance and cell migration phenotypes and come with a large body of mutational and functional data. To study UNC-6–UNC-5 interactions and possible roles for heparin in this interaction, we established a yeast display system for UNC-5 and show that the binding between UNC-5 and UNC-6 is strongly enhanced when heparin is added. Through directed evolution of UNC-5 displayed on yeast, we have identified and engineered the UNC-5 interface used for heparin binding, and we confirmed heparin binding to this surface with a crystal structure of a complex formed between UNC-5 and a short form of heparin. Addition of heparin allows for the formation of UNC-6–UNC-5 ectodomain complexes, as well as the complete repulsive complex, including UNC-6 and the ectodomains of UNC-5 and UNC-40/DCC in a large oligomeric entity, with a surprisingly rigid and globular structure. Finally, we have shown that abolition of heparin affinity of UNC-5 in *C. elegans* results in cell migration defects associated with loss of function of *unc-5* and *unc-6*, indicating mechanistic dependence of Netrin:UNC-5-mediated guidance on heparan sulfate. Our results provide the first detailed glimpses into repulsive Netrin complexes and may be crucial in explaining chemo- and haptotactic responses to Netrin sources.

## RESULTS

### High-affinity UNC-6–UNC-5 binding depends on heparin

The interactions of Netrins with their attractive receptors, DCC and Neogenin, have been heavily characterized using structural and mutational studies (Finci et al., 2014; Xu et al., 2014). However, structural data for Netrin interactions with its repulsive receptor, UNC-5, remain sparse, likely due to the low affinity for this ligand-receptor pair. To study UNC-6–UNC-5 interactions, we established the use of yeast surface display as a platform capable of measuring and engineering this interaction, which further enabled us to also investigate the effects of additional molecular players, most importantly glycosaminoglycans. Briefly, we displayed on yeast cells the ectodomain of *C. elegans* UNC-5, including its two immunoglobulin (IG) and two Thrombospondin type I (Tsp1) domains, by way of a fusion to the yeast cell wall protein, Aga2 (**Figure 1A**). UNC-5 is expressed on yeast cells as evident from antibody staining of the C-terminal myc tag. To measure Netrin binding to UNC-5-expressing cells, we expressed and purified a soluble construct of *C. elegans* UNC-6/Netrin lacking its C-terminal NTR domain (UNC-6ΔC), since the NTR domain is known to cause aggregation (Keino-Masu et al., 1996). We created a yeast-staining reagent by biotinylating UNC-6ΔC (**Figure S1A**) and incubating it with Alexa Fluor 647-coupled streptavidin. In the absence of heparin, UNC-6ΔC binds very weakly to UNC-5-displaying yeast, as measured with flow cytometry by the weak fluorescence shift relative to the negative control yeast (**Figure 1B red**). To test if the interaction can be improved by glycosaminoglycans, we added heparin (from porcine intestinal mucosa, ~15 kDa) to the UNC-6-labeling experiments of UNC-5-expressing yeast, which showed strong binding (right shift of the fluorescence histogram) with an apparent *K*_D_ of 35 nM in the presence of 5 μg/ml (333 nM) heparin (**Figures 1A right, 1B green, and S1C**). This demonstrates that strong UNC-6–UNC-5 binding may require heparin, likely in the form of a Heparan Sulfate proteoglycan on the growth cone or the substrate tissue *in vivo*.

**Figure 1.**
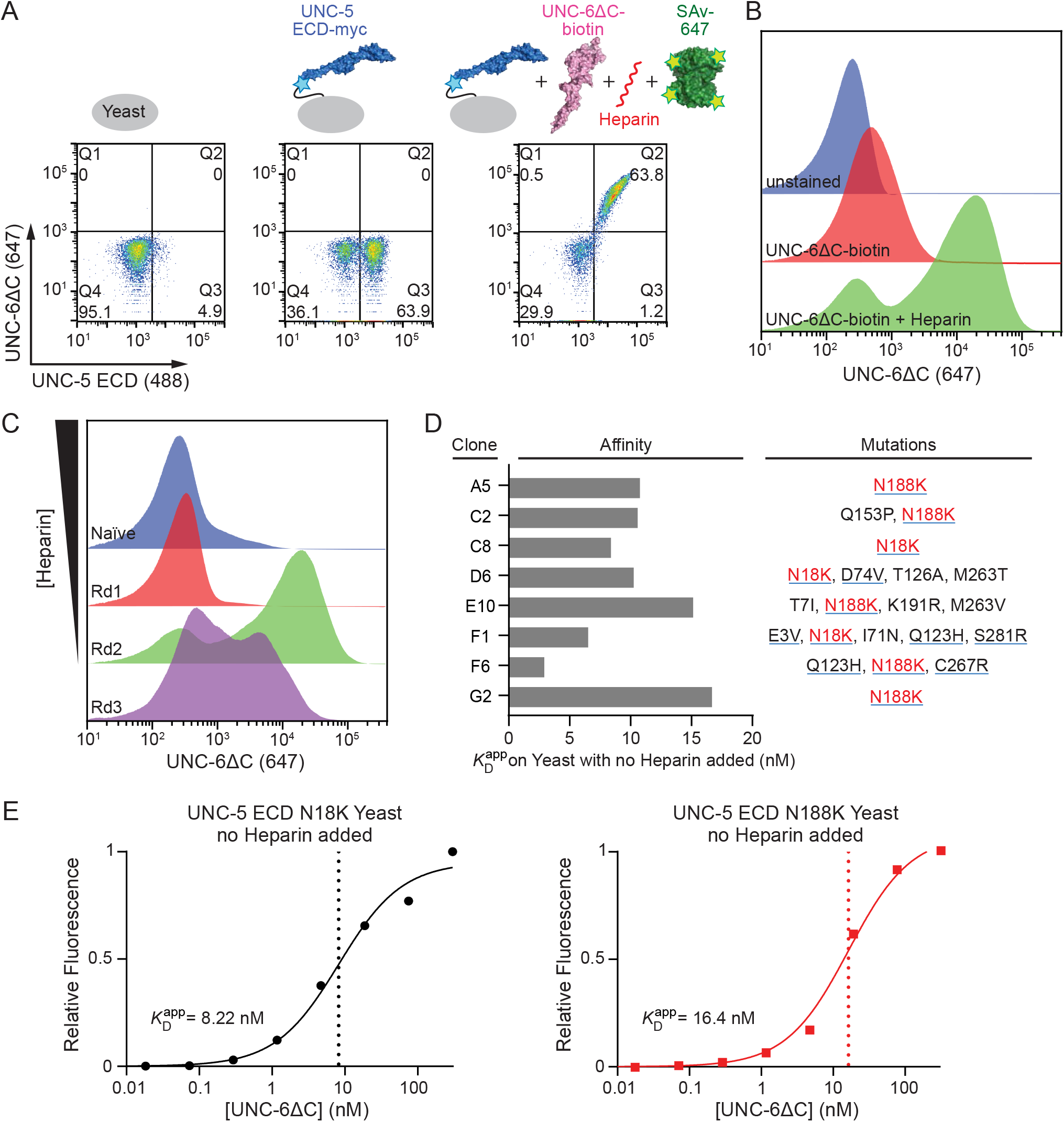
Directed Evolution of UNC-5 ectodomain. **A.** UNC-5 ECD displays on yeast, as detected by anti-myc staining (Alexa 488), and strongly interacts with biotinylated UNC-6ΔC monomers bound to Alexa 647-coupled Streptavidin in the presence of heparin. (*Left*) Unstained cells, (*Middle*) Cells stained with anti-myc antibody, (*Right*) Cells double-stained with anti-myc and UNC-6ΔC–streptavidin with 5 μg/ml heparin added. **B.** Histograms for Alexa 647 fluorescence of unstained cells (blue), UNC-6ΔC-stained cells (red), and UNC-6ΔC-stained cells in the presence of 5 μg/ml heparin (green). **C.** Histograms of bulk UNC-6ΔC labeling of the yeast library during selections, where heparin concentration was decreased. The final, third round of selection contained no added heparin. **D.** Apparent UNC-6ΔC-binding affinities for the selected UNC-5 variants on yeast. All binding isotherms are shown in Figure S1A. Mutations increasing net positive charge on UNC-5 ECD are underlined blue; no mutation in the selected colonies added net negative charge. Data were collected in the absence of added heparin. **E.** Binding isotherms of two single-site mutants of UNC-5 ECD, N18K and N188K, for UNC-6ΔC in the absence of heparin.

It should be noted that even without its C-terminal domain, UNC-6 remains a highly charged protein that aggregates, as recently reported by Krahn et al. (2019), or non-specifically interacts with chromatography resin. We discovered that UNC-6ΔC can be purified and stored in higher salt to improve its behavior in size-exclusion chromatography (SEC) runs, best in the presence of 150 mM NaCl supplemented with 100 mM MgSO_4_ (**Figure S1B**).

### Directed evolution of heparin dependence of UNC-6–UNC-5 binding

We hypothesized that we can select for UNC-5 mutants with a lower dependence on heparin for UNC-6 binding using directed evolution. We created an UNC-5 ectodomain display library using random error-prone PCR and selected for UNC-6ΔC binding while decreasing heparin concentrations in the staining solution (**Figure 1C**). The third and final round of selections was performed in the absence of added heparin and therefore shows a slight reduction in binding compared to the previous round. We were able to evolve high-affinity UNC-6 binding as seen in titrations on yeast (**Figures 1D and S1C**). Sequencing of eight clones from the third round of selection shows that every clone included either an N18K or an N188K mutation (**Figure 1D**). As these mutations increase net positive charge on UNC-5, we predict that we may have inadvertently increased UNC-5 affinity to heparin, in addition to strengthening binding to UNC-6. Since we could observe strong UNC-6 binding with these mutants even in the absence of added heparin, binding might be supported by an endogenous, unknown GAG-like molecule on the yeast surface.

To confirm that the two mutations identified are the cause for increased UNC-6ΔC binding in yeast display experiments, we performed titrations of UNC-6ΔC binding to UNC-5 N18K/N188K ectodomain on yeast in the absence of added heparin. (**Figure 1E**). Apparent binding affinities of UNC-6ΔC to UNC-5 N18K and N188K-expressing yeast were relatively strong and in the nanomolar range, 8.2 and 16.4 nM, respectively.

### UNC-5 binds heparin via a positively charged patch at the boundary of its IG1 and IG2 domains

To gain more insights into UNC-5 interactions with heparin and/or UNC-6, we set out to determine the structure of the *C. elegans* UNC-5 ectodomain. Crystallization was successful for a construct containing the two IG domains, which included both N18 (IG1) and N188 (IG2) identified in directed evolution experiments. Using these crystals, we determined the structure of the two-IG domain UNC-5 construct to 2.9 Å resolution (**Figure 2A**). The structures of the immunoglobulin domains show no major differences to previously determined human Unc5A, rat Unc5D (Jackson et al., 2016; Seiradake et al., 2014) and mouse Unc5B crystal structures (Stetefeld et al., 2020), including a buried disulfide bond linking UNC-5 Cys37 (C strand) and Cys84 (F strand) in IG1 in addition to the canonical disulfide bond found in IG domains, and a short α-helix connecting E and F strands in IG1. Similarly, the IG2 domain is N-glycosylated at a remarkably conserved site, Asn178 (F strand), also observed in the rat Unc5D and mouse Unc5B structures; no glycosylation on IG1 is observed. We also had the opportunity to compare flexibility at the IG1-IG2 boundary: An alignment of our UNC-5 IG1+2 with three other structures show wide variability of the angle between the IG1 and IG2 domains (**Figure S2A**). Flexibility at this position may be necessary for recognition by heparan sulfate as well as protein ligands, since our directed evolution results indicate this as a site of heparan sulfate binding, and this is also the binding site for Latrophilins (Jackson et al., 2016).

**Figure 2.**
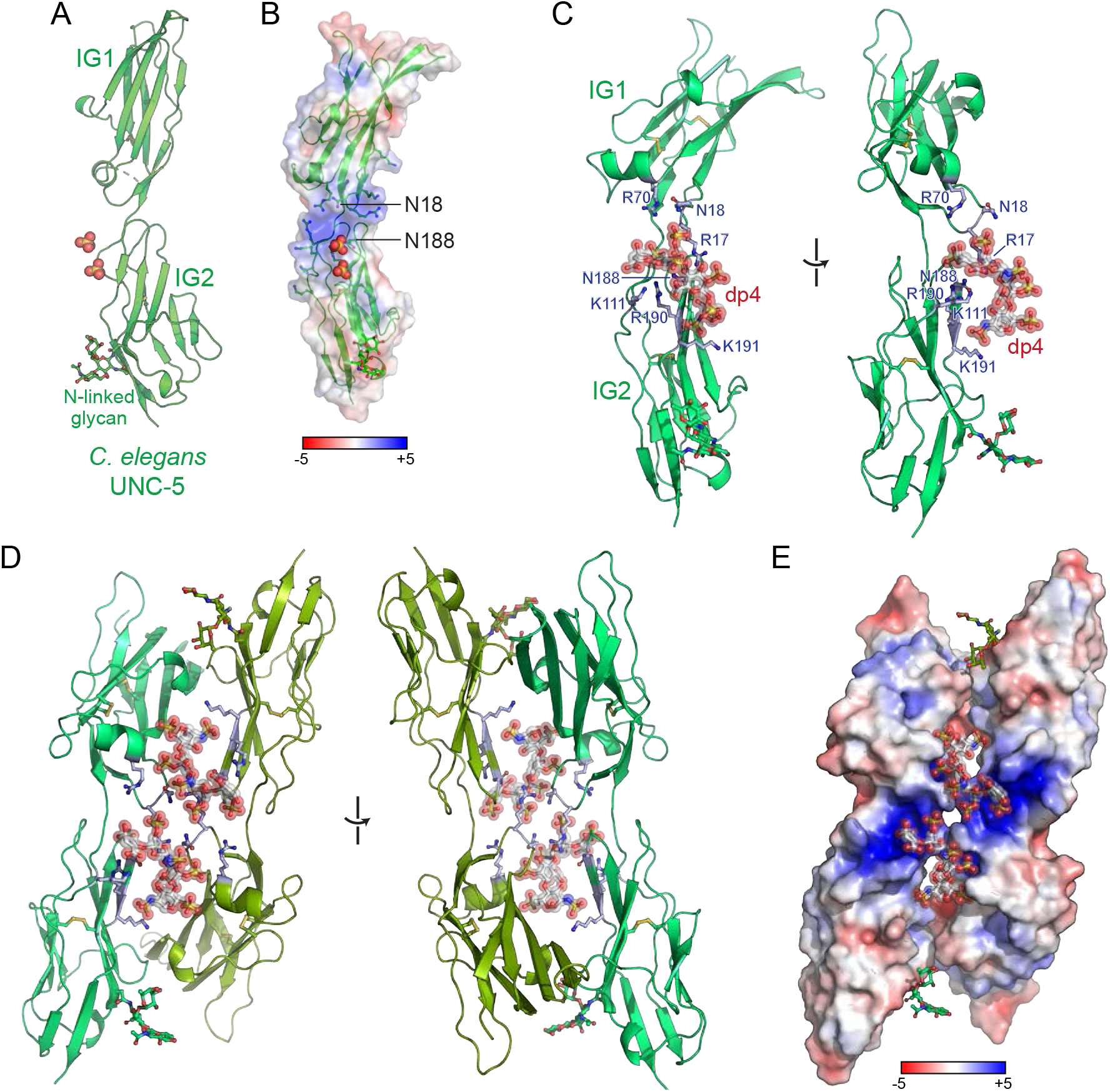
Crystal structures of UNC-5 IG1-2 highlight a positively charged surface used for heparan sulfate binding. **A.** UNC-5 IG1-2 structure with putative sulfates bound near the IG1-IG2 boundary. The conserved N-linked glycan at N178 is shown in ball-and-stick representation. **B.** Electrostatic surface potential for the UNC-5 IG1-2 structure, showing a positively charged surface at the IG1-IG2 boundary. The range of electrostatic potential used to color the surface is from −5 to +5 *kT*/*e*. The two Asn residues identified in directed evolution experiments are labeled. **C.** Crystal structure of the UNC-5 IG1-2 domains bound to the heparin tetrasaccharide (dp4), one alternate conformation displayed. **D.** The asymmetric unit of the UNC-5+dp4 crystal structure, showing a possible dimer created by a dp4 molecule, built as two alternate conformations into density (**Figure S2**). The two alternate conformations overlap with two alternates of a dp4 molecule from the neighboring asymmetric unit, creating an intimate four-molecule UNC-5 entity, as seen in Figure S2D. **E.** Surface potential of the dimer shows a contiguous positive surface, which is used for binding to heparan sulfate.

Interestingly, there was unexplained electron density surrounding the IG1-IG2 boundary (**Figure S2B**). Based on the presence of ammonium and lithium sulfate in the crystallization solution and the chemistry around the unexplained density, we believe that there may be two or more fully or partially occupied sulfate ions at this site. Surface electrostatic potential of our UNC-5 structure shows an intensely positive region at this domain boundary created by multiple arginines and lysines (**Figure 2B**). Furthermore, the two N→K mutations selected in our directed evolution experiments fall in this region interspersed between the positively charged arginines and lysines.

Overall, this site is a strong candidate for heparan sulfate binding.

We next crystallized and determined the structure of UNC-5 IG1+2 domains in the presence of a short heparin oligomer limited to four saccharide units, dp4, in a different crystal form. We observed strong extra density resembling an oligosaccharide oligomer in the same region, surrounded by Arginines 17, 70 and 190, Lysines 111 and 191 and Asparagines 18 and 188 (**Figure 2C**).

The UNC-5–heparin-dp4 crystals have an UNC-5 dimer in the asymmetric unit created by 1210 Å^2^ of buried surface area, which appears to be held together partly by heparin interacting with both chains (**Figure 2D**). As there is some ambiguity in the exact orientation of the dp4 ligand and based on electron density, we chose to model dp4 in two alternate conformations in equivalent positions within this dimer, which then overlap with their alternate conformations from the neighboring asymmetric unit. An intimate tetramer, also held together by and surrounding heparin molecules is observed upon investigation of neighboring asymmetric units, where protein-protein contacts bury 1310 Å^2^ and 1120 Å^2^ between UNC-5 chains related by crystallographic symmetry (**Figures S2C and S2D**). While these molecular interfaces may be physiological in the context of a heparin-mediated UNC-5 or UNC-6–UNC-5 oligomer, we do not yet have evidence to support such a model.

### The IG–heparin interaction observed in the UNC-5–heparin complex may be a common feature of IG-superfamily receptors

Due to its sulfation and negative charge, heparin binding strongly favors positively charged surfaces on proteins. As a strongly positively charged protein, UNC-6 (pI = 8.7) binding to heparin is not surprising. However, UNC-5 ectodomain has a pI of 5.7, and a net predicted charge of –6 at pH 7. The heparin–UNC-5 interactions are a result of a local charge at the IG1-IG2 boundary as observed in our crystal structure (**Figure 2E**). The positively charged amino acids that create the HS-binding site in nematode UNC-5 are conserved across all major bilaterian taxa (**Figures 3A and S3**) and make up the major large patch of conserved surface in UNC-5 ectodomains (**Figure 3B**).

**Figure 3.**
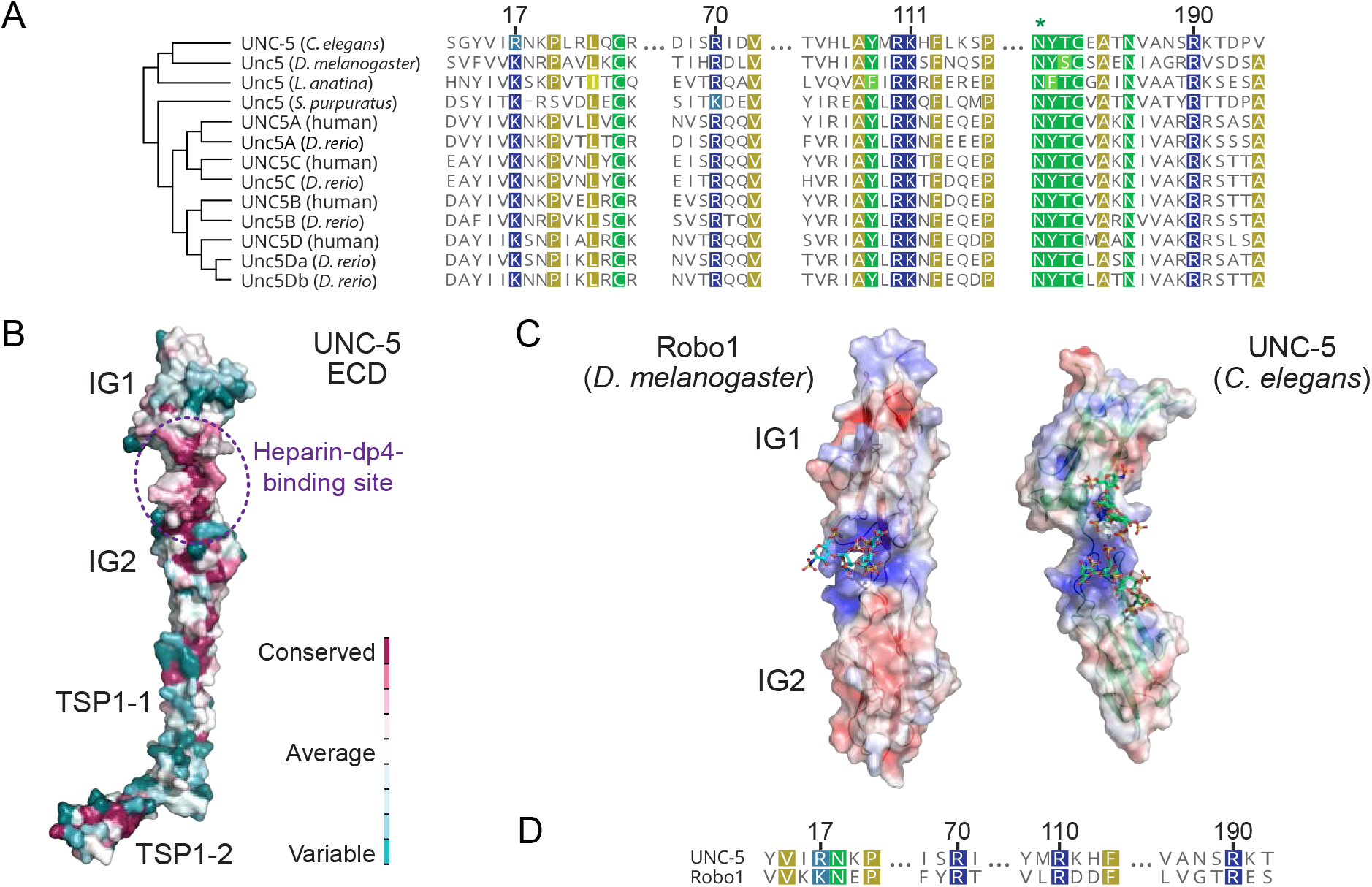
The positively charged patch at the UNC-5 IG1-IG2 boundary is conserved. **A.** Alignment of protostome and deuterostome UNC-5 sequences show the positive charge at the IG1-IG2 boundary to be highly conserved. Sequence numbering above follows the *C. elegans* UNC-5 sequence. **B.** Surface conservation analysis by ConSurf of the full ectodomain of UNC-5 (using the AlphaFold model) shows the IG1-IG2 boundary to be largest conserved surface patch. **C.** Side-by-side views of the C. elegans UNC-5 IG1-2+dp4 structure and fruit fly Robo1 IG1-2+dp8 structure (PDB ID: 2VRA; only a tetrasaccharide is visible), with electrostatic surface potentials. [Check the details here] **D.** Sequence alignment of the UNC-5 and Robo1 IG1-2 domain contacts, showing the positively charged patch at the IG1-IG2 boundary are common to both receptors.

Since heparin has been implicated in binding to several neuronal adhesion and signaling receptors, including other immunoglobulin superfamily proteins, we looked for structural clues in other IG-type receptors with heparin binding. Robo receptors, which are also repulsive guidance receptors, bind heparin using the same surface at their IG1-IG2 boundary (**Figures 3C and 3D**) (Fukuhara et al., 2008), suggesting a conserved or convergent site for heparin interactions among immunoglobulin superfamily (IgSF) neuronal receptors for heparin binding, although the binding poses of the heparin molecules may be different.

### Biochemical characterization of UNC-5 mutants selected by directed evolution and rational design

Through directed evolution and structural biology, we have identified a positively charged surface at the UNC-5 IG1-IG2 boundary, which interacts with heparin, and can be mutated to increase UNC-6 affinity. To confirm and extend these findings using orthogonal methods, we performed surface plasmon resonance experiments with UNC-5 mutants for binding heparin and/or UNC-6. First, we showed that UNC-6–UNC-5 binding is weak with a *K*_D_ of 46 μM as measured by biotinylated UNC-6ΔC captured on an SPR chip (**Figure 4A**). The N18K and N188K mutations on UNC-5 improve binding to UNC-6 progressively: UNC-5 N18K and N188K bind UNC-6ΔC with an affinity 2-fold and 3-fold improved over wild-type (WT) UNC-5, respectively, confirming the results of our directed evolution experiments (**Figures 4A and 4C**). Next, we showed that UNC-5 and UNC-6 both bind heparin separately (**Figures 4D and 4F**). UNC-6ΔC binding to heparin is strong – with a *K*D of 39 nM, while UNC-5 affinity for heparin is weaker (*K*_D_ = 26 μM). We measured UNC-6ΔC–heparin affinity with UNC-6 captured on the chip, since UNC-6ΔC can best be used with a buffer containing MgSO_4_ (**Figure S1B**), which would interfere with heparin binding. UNC-5–heparin binding, on the other hand, could be measured using biotinylated heparin captured on an SPR chip, and soluble UNC-5 as analyte. The UNC-5 N18K and N188K mutations, which are at the heparin binding site according to our crystal structure, improve heparin binding as expected, with the double N→K UNC-5 mutant binding heparin with a 16-fold increase in affinity (*K*_D_ = 1.7 μM) (**Figures 4D and 4E**).

**Figure 4.**
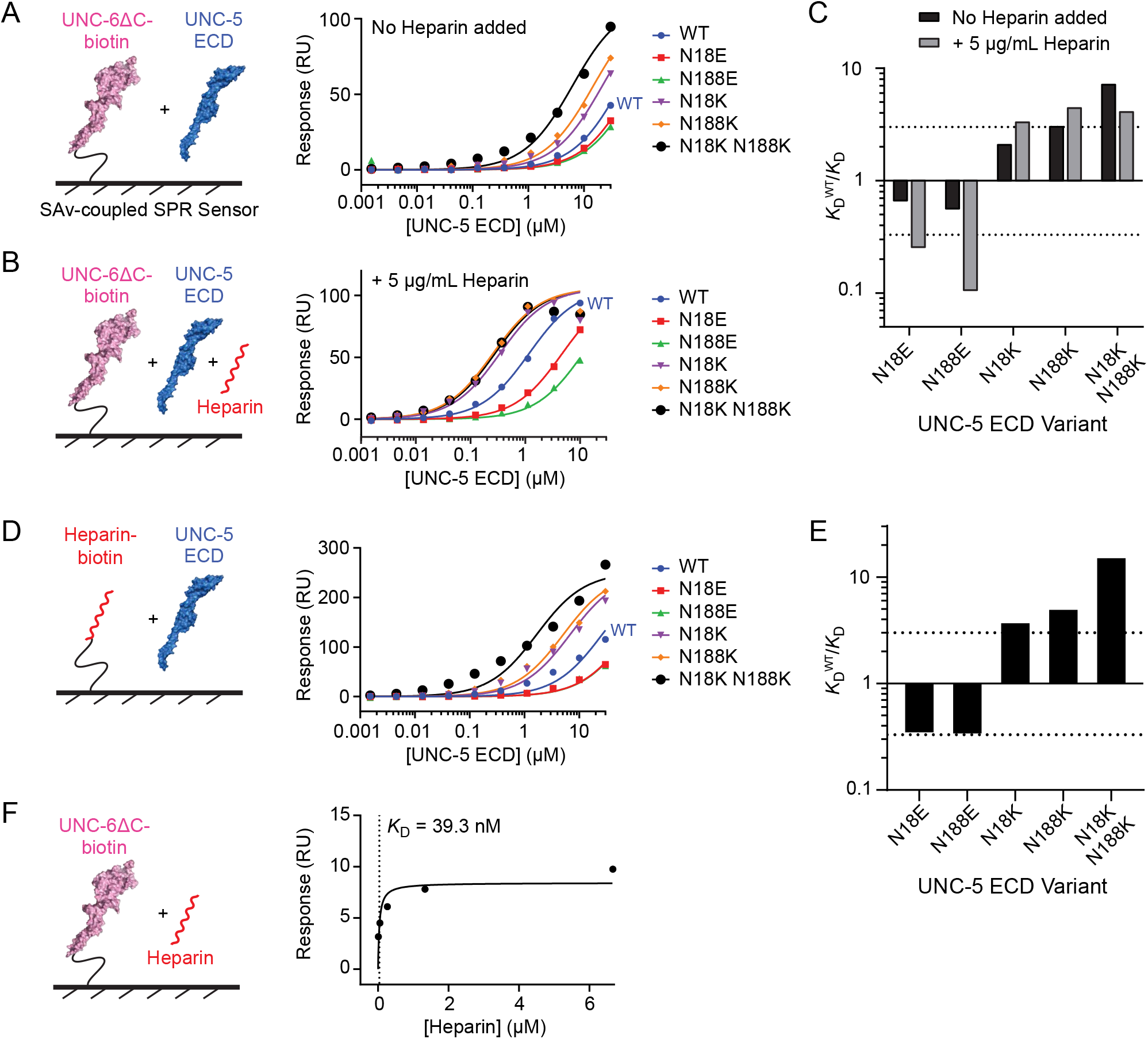
UNC-5 mutations at N18 and N188 modulate UNC-5 affinity to UNC-6 and heparin. **A, B.** Binding isotherms for SPR experiments testing the interaction of biotinylated WT UNC-6ΔC with UNC-5 ECD WT and mutants without added heparin (A), and in the presence of 5 μg/ml heparin (B). **C.** Relative binding affinities to UNC-6ΔC of UNC-5 ECD mutants compared to WT without added heparin (black) and in the presence of 5 μg/ml heparin (gray). Dotted line represents a 3-fold difference. **D.** Binding isotherms for SPR experiments testing the interaction of biotinylated heparin with UNC-5 ECD WT and mutants. **E.** Relative binding affinities to heparin of UNC-5 ECD mutants. Dotted line represents a 3-fold difference. **F.** Binding isotherm for SPR experiment testing the interaction of biotinylated WT UNC-6ΔC with heparin. All sensograms are shown in Figure S4.

We observed in our yeast display experiments that the UNC-5–UNC-6 affinity is significantly improved in the presence of heparin. We also confirmed these results with SPR: UNC-5 binds UNC-6ΔC with a *K*_D_ of 1.2 μM in the presence of 5 μg/mL (333 nM) heparin, an improvement of 39-fold compared to UNC-5–UNC-6ΔC binding in the absence of heparin (**Figures 4A and 4B**). At this heparin concentration, we expect that nearly all UNC-6 molecules are in complex with heparin, but WT UNC-5 may be mostly unoccupied with heparin. As the two Asn’s at the IG1-IG2 boundary can be mutated to positively charged Lys to improve UNC-5 affinity for both heparin and UNC-6, we expect that UNC-5 binding to UNC-6 in the presence of heparin should also be improved with these mutations: UNC-5 ectodomains with N18K and N188K mutations progressively bind UNC-6 more strongly in the presence of heparin, with a 4-fold improvement for the double mutant (**Figures 4B and 4C**).

Based on our yeast display and structural work, we were also able to create partial loss-of-function UNC-5 mutants by mutating the two Asn residues to Glu’s. UNC-5 ectodomain with either N18E or N188E mutations show weaker binding to heparin (**Figures 4D and 4E**), and weaker binding to UNC-6 in the presence or absence of heparin (**Figures 4A-C**), with an affinity 4 to 10-fold worse than the wild-type UNC-6–UNC-5 interaction in the presence of heparin, and 1.6 to 1.8-fold worse in the absence of heparin.

While the N→E mutants of UNC-5 have lower affinity for UNC-6 as expected, binding is not entirely abolished due to the presence of extensive positive charge at the IG1-IG2 boundary of the UNC-5 ectodomain. Based on the crystal structure of the UNC-5–heparin complex, we designed four mutants of UNC-5, reversing charge at the heparin-binding site. Mutants contained a subset or all of the following amino acid substitutions: R17E, R70E, K11E, R190E, and R191E. All of the mutants showed a drastic reduction; e.g., UNC-5 ectodomains with the triple mutation R17E K111E R190E lost heparin-binding affinity 51-fold compared to WT (**Figures 5A and 5B**). Binding to UNC-6, in the absence or presence of heparin, is also affected with all mutants, but to a much greater degree when heparin is present (**Figures 5C-E**): For example, UNC-5 R17E K111E R190E binds UNC-6ΔC in the presence of 5 μg/mL heparin with a ~187 μM dissociation constant. At this heparin concentration, we expect UNC-6 to be mostly heparin bound, but the UNC-5 triple mutant to be free of heparin. These results imply that UNC-5 binding to heparin strongly facilitates UNC-6 binding, and that there are additional weak contacts between UNC-5 and UNC-6 that are not heparin dependent.

**Figure 5.**
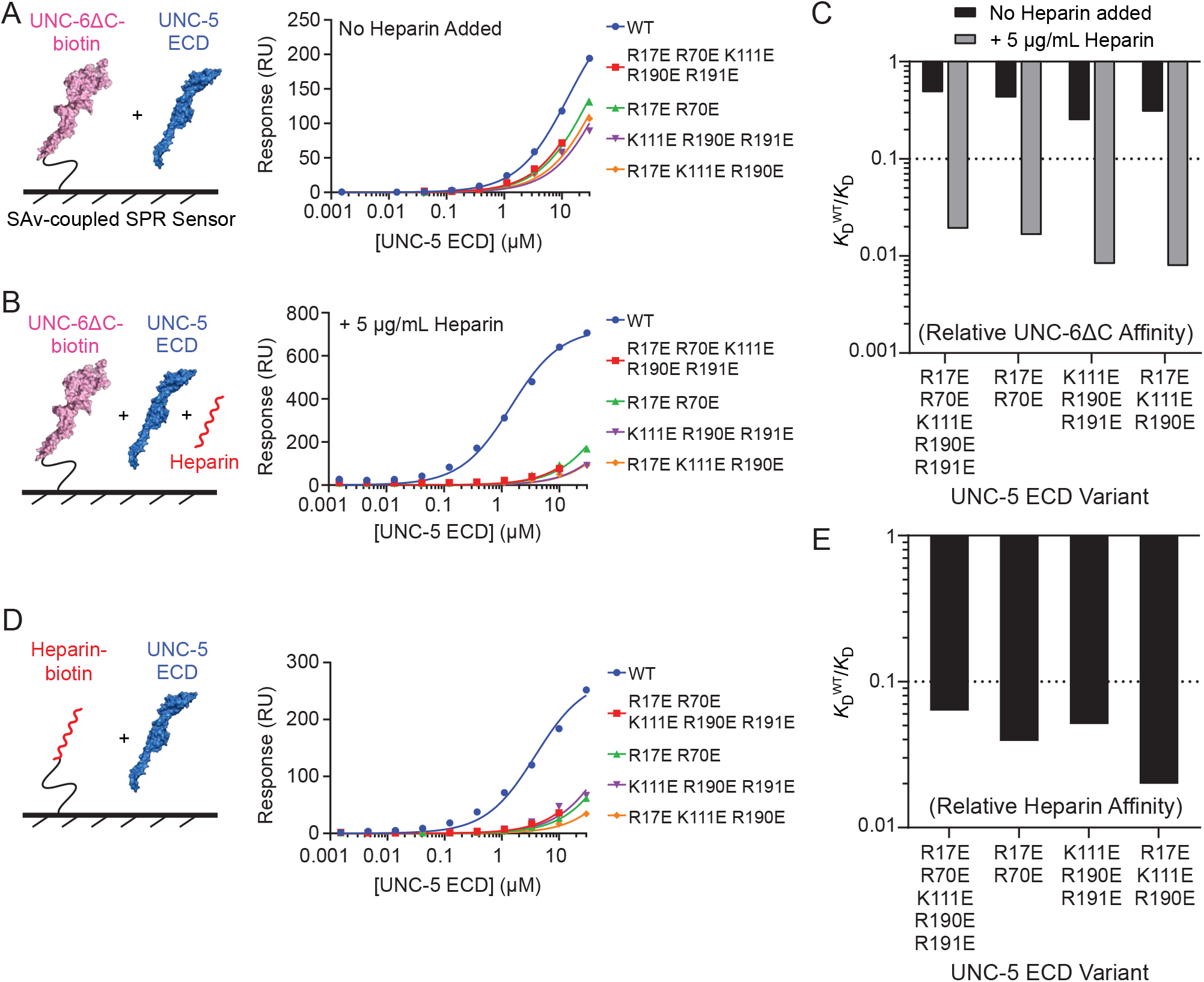
Loss of heparin binding by UNC-5 accompanies loss of UNC-6 binding. **A.** Binding isotherms for SPR experiments testing the interaction of biotinylated heparin with UNC-5 ECD WT and mutants. UNC-5 ECD mutants in Figure 5 contain a subset or all of the following amino acid substitutions designed to disrupt heparin binding: R17E, R70E, K11E, R190E, and R191E. **B.** Relative binding affinities to heparin of UNC-5 ECD mutants. Dotted line represents a 10-fold difference. All sensograms are show in Figure S5. **C, D.** Binding isotherms for SPR experiments testing the interaction of biotinylated WT UNC-6ΔC with UNC-5 ECD WT and mutants without heparin (A) and in the presence of 5 μg/ml heparin (B). **E.** Relative binding affinities to UNC-6ΔC of UNC-5 ECD mutants compared to WT without added heparin (black) and in the presence of 5 μg/ml heparin (gray). Dotted line represents a 10-fold difference.

### Heparin binding creates a stable, oligomeric repulsive Netrin-Receptor complex

Since heparin can bind both UNC-6/Netrin and UNC-5, and increase their affinity to each other, we next asked the question if we can reconstitute a stable extracellular complex of UNC-6 and UNC-5. Over size-exclusion runs, we could not see a stable UNC-6ΔC–UNC-5 ECD complex without heparin (**Figure 6A**), which can be explained by their low affinity in the absence of heparin (**Figure 4A**). When heparin was added at a 1:1:1.5 UNC-6:UNC-5:heparin molar ratio, a stable complex eluting at a volume corresponding to a ≥1 MDa complex was observed (**Figure 6B**). SDS-PAGE analysis of the complex peak show a complex including a 1:1 molar ratio of UNC-6:UNC-5 (**Figures 6B and S6A**).

**Figure 6.**
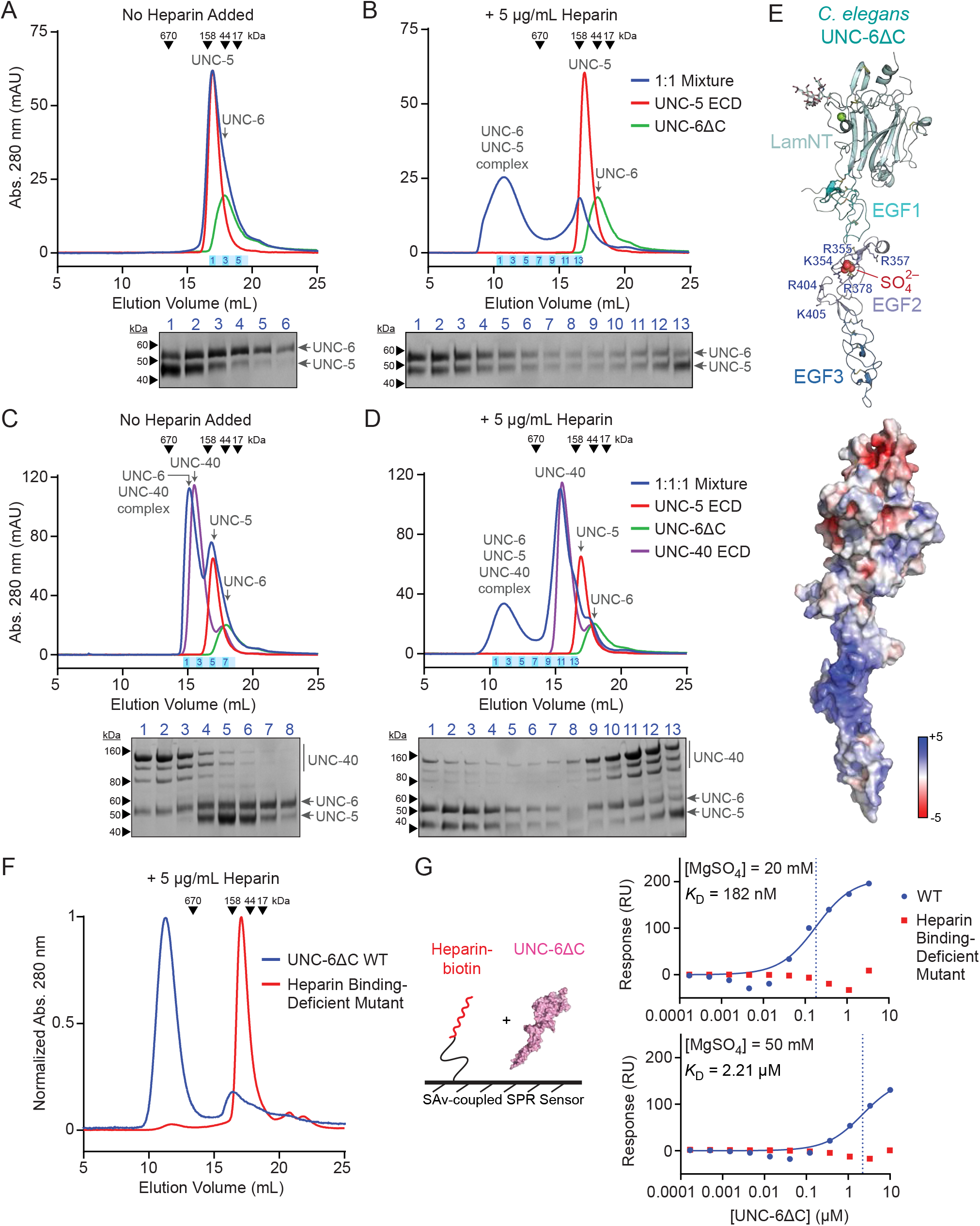
UNC-5 and UNC-6 form a large oligomeric complex in the presence of heparin, with the ability to incorporate UNC-40. **A,B.** Size-exclusion chromatography of UNC-5 ECD and UNC-6ΔC. UNC-5 ECD (red), UNC-6ΔC (green), and a 1:1 molar ratio mixture (blue) were injected without added heparin (A) and with 5 μg/ml heparin (B) on a column equilibrated in HBS-MS and no additional heparin. Chromatograms in A-D and F show elution profiles measured by absorbance at 280 nm with a pathlength of 0.2 cm, using a Superose 6 Increase 10/300 column along with SDS-polyacrylamide gels of eluted fractions from the runs with mixtures (blue). Coomassie-stained bands are quantified in Figure S6A. **C,D.** Size-exclusion chromatography of UNC-5 ECD, UNC-6ΔC, and UNC-40 ECD. UNC-5 ECD (red), UNC-6ΔC (green), UNC-40 ECD (purple), and a 1:1:1 mixture (molar ratio, blue) were injected without heparin (C) and with 5 μg/ml heparin added (D) on a Superose 6 10/300 column equilibrated with HBS-MS. Coomassie-stained SDS-polyacrylamide gels for the mixture runs (blue) are placed under the chromatograms, and are quantified in Figure S6A. **E.** Crystal structure (top) and electrostatic surface potential (bottom) of UNC-6ΔC. The range of electrostatic potential used to color the surface is from −5 to +5 *kT/e*. **F.** Size-exclusion chromatography of UNC-6ΔC WT (blue) and the heparin binding-deficient mutant (red) mixed with 5 μg/ml heparin. **G.** Binding isotherms for SPR experiments testing the interaction of biotinylated heparin with UNC-6ΔC WT (blue) and the heparin binding-deficient mutant (red) at two MgSO_4_ concentrations to counter non-specific binding.

It has been suggested that heparin binding might bias the binding of Netrins towards specific receptors (Meijers et al., 2020). If that is the case, we may expect heparin to break the otherwise stable UNC-6 complex with UNC-40/DCC. We confirmed that UNC-6ΔC and UNC-40 ECD can make a stable complex containing an equal molar ratio of UNC-6 and UNC-40 (**Figure 6C and S6A**). When UNC-6ΔC, UNC-5 and UNC-40 ECDs are mixed in the presence of heparin and applied to an SEC column, we observed a large oligomeric peak containing the UNC-6:UNC-5:UNC-40 complex with a largely substoichiometric UNC-40 content, and excess free UNC-40. We have estimated a complex content of approximately 5:5:1 UNC-6:UNC-5:UNC-40 based on Coomassie staining (**Figures 6D and S6A**). This implies that heparin and/or UNC-5 may partly compete for UNC-40 binding, and that UNC-40 may contribute to repulsive signaling as part of a Netrin–UNC-5 complex at a substoichiometric ratio. Removing heparin from this SEC run resulted in the loss of the MDa-sized complex peak (**Figure 6C**). To our knowledge, this is the first demonstration of a Netrin–UNC-5–DCC–heparin complex purified at a preparative scale, even though UNC-40/DCC is binding at a substoichiometric ratio.

### Heparin-mediated UNC-6 oligomerization

We next investigated how the UNC-6–UNC-5 complex could oligomerize and hypothesized that this could be due to heparin-mediated oligomerization of UNC-6. To gain more insights into the UNC-6 structure, we were able to determine the structure of UNC-6ΔC, but not with heparin bound (**Figure 6C**). However, we observed a highly ordered sulfate ion at a highly positively charged site on the second EGF domain of UNC-6 (**Figure 6D**), close to where Netrin–UNC-5 binding was proposed based on antibody-blocking experiments (Grandin et al., 2016). The positive charge at this site is conserved in the monophyletic Netrin-1/3/5 family of proteins, but not Netrins that separately evolved and are not known to mediate axon guidance (Netrin-4 and Netrin-G families) (**Figure S6B**) (Finci et al., 2014; Leclère and Rentzsch, 2012). Finci et al. (2014) has also observed putative sulfate ions in their Netrin-1–DCC crystal structure at the second EGF domain, and proposed the positively charged surface on the EGF2 domain as a potential heparin-binding site.

To test if the positively charged surface on EGF2 is the heparin-binding site outside the C-terminal NTR domain, we created an UNC-6ΔC mutant where K354, R355, R357, R378, R404, and K405 were mutated to glutamate. When WT UNC-6ΔC is run on an SEC column after being mixed with heparin at a 1:1.5 molar ratio, it runs as a large oligomer (**Figure 6F**). The engineered UNC-6ΔC mutant (“Heparin Binding-Deficient Mutant” in **Figure 6**), however, does not oligomerize when mixed with heparin. We also performed SPR experiments to measure heparin affinity for WT and mutant UNC-6ΔC, where biotinylated heparin was captured on an SPR chip, and soluble UNC-6ΔC was used as an analyte at a variety of MgSO_4_ concentrations to counter non-specific binding (**Figure 6G**). The heparin binding-deficient mutant showed no binding, while we observed nM to μM affinity depending on the MgSO_4_ concentration: As expected, higher [MgSO_4_] results in weaker binding, while lower [MgSO_4_] are in the nM range, similar to our measurements in Figure 4F, where binding could be measured in no MgSO_4_. As heparin can still bind and oligomerize WT UNC-6ΔC even in the presence of 100 mM SO_4_^2−^ in the column running buffer, UNC-6ΔC is clearly a potent heparin-binding protein. Our results show that the positively charged site on the EGF2 domain is a heparin-binding site, which also helps oligomerize UNC-6 in the presence of heparin. Given these data, the most likely explanation for the formation of the UNC-6–UNC-5–heparin oligomers is heparin-mediated UNC-6 oligomerization.

### Biophysical properties of the heparin-mediated UNC-6–UNC-5 oligomers

An important question relevant to UNC-5 signaling is whether the heparin and UNC-6-mediated UNC-5 oligomers are an ordered and well-defined molecular species, or these oligomers are simply a result of UNC-6 and UNC-5 decorating the long, linear heparin chains. We studied this in solution using small-angle x-ray scattering (SAXS), as a mixture of UNC-6ΔC, UNC-5 ECD and heparin were injected on an SEC column, and SAXS data was collected as the complex eluted (**Figure S6D**). The pair distribution (*P*(*r*)) plot and the molecular shape analysis both indicate a roughly globular complex, and the dimensionless Kratky plot shows a folded and highly rigid molecule (**Figures S6E-G**). Given inherent flexibility of the heparin chain and the multi-domain proteins involved, including the flexible UNC-5 ectodomain (Figure S2A), this is unexpected, and indicates that the UNC-6–UNC-5–heparin complex has a stable, ordered geometry and well-defined structure, which may be important for a signaling-competent conformation on the cell surface. Addition of UNC-40 to the SEC-SAXS sample, results in little difference in the scattering data (**Figures S6D-F**), where the substoichiometric UNC-40 ECD may not significantly affect the shape and structure of the complex but may contribute to signaling via its intracellular signaling motifs.

### Heparin-binding activity of UNC-5 is necessary for UNC-6-controlled dorsal migration of cells during development

To reveal if a functional Netrin–UNC-5 complex depends on the ability of UNC-5 to interact with heparin, we engineered one of our heparin binding-deficient UNC-5 mutants, R17E R70E, in *C. elegans*. Early studies have identified a requirement for UNC-6–UNC-5 signaling in dorsal guidance of both motor axons and distal tip cells. The mobile distal tip cell is the leading cell of the developing gonad that migrates along the ventral side of the animal before turning dorsally in hermaphrodites. These cells continue to migrate to the dorsal-most region of the animal before turning again to continue migration along the anterior-posterior axis (**Figure 7A**) (Hedgecock et al., 1987). The migration route of the distal tip cells establishes the U-shaped morphology of the gonads in the adult hermaphrodite (Kimble and White, 1981). These migrations require both *unc-6* and *unc-5* (Hedgecock et al., 1990), which is consistent with a repulsive action against a ventral source of UNC-6/Netrin (Hamelin et al., 1993; Wadsworth et al., 1996). Additionally, the HSPG LON-2/Glypican modifies UNC-6–UNC-5 signaling in the migrating distal tip cells (Blanchette et al., 2015). We observed that in *unc-5(R17E R70E)* animals the majority of distal tip cells either do not turn dorsally or fail to migrate the full length of the dorsal-ventral body axis (**Figures 7B and 7C**). We compared the phenotype from these animals with mutant strains *unc-6(ev400)* and *unc-5(e53)*, previously identified as null mutants for dorsal migration-related functions (Hedgecock et al., 1990). The failed distal tip cell migrations of *unc-5(R17E R70E)* mutants are indistinguishable from the defects observed in null *unc-6* and *unc-5* mutant animals (**Figure 7C**). These results indicate that the Netrin-dependent UNC-5 function requires the heparin-binding activity of UNC-5, as this is likely necessary for a stable, oligomeric and functional UNC-6–UNC-5 complex.

**Figure 7.**
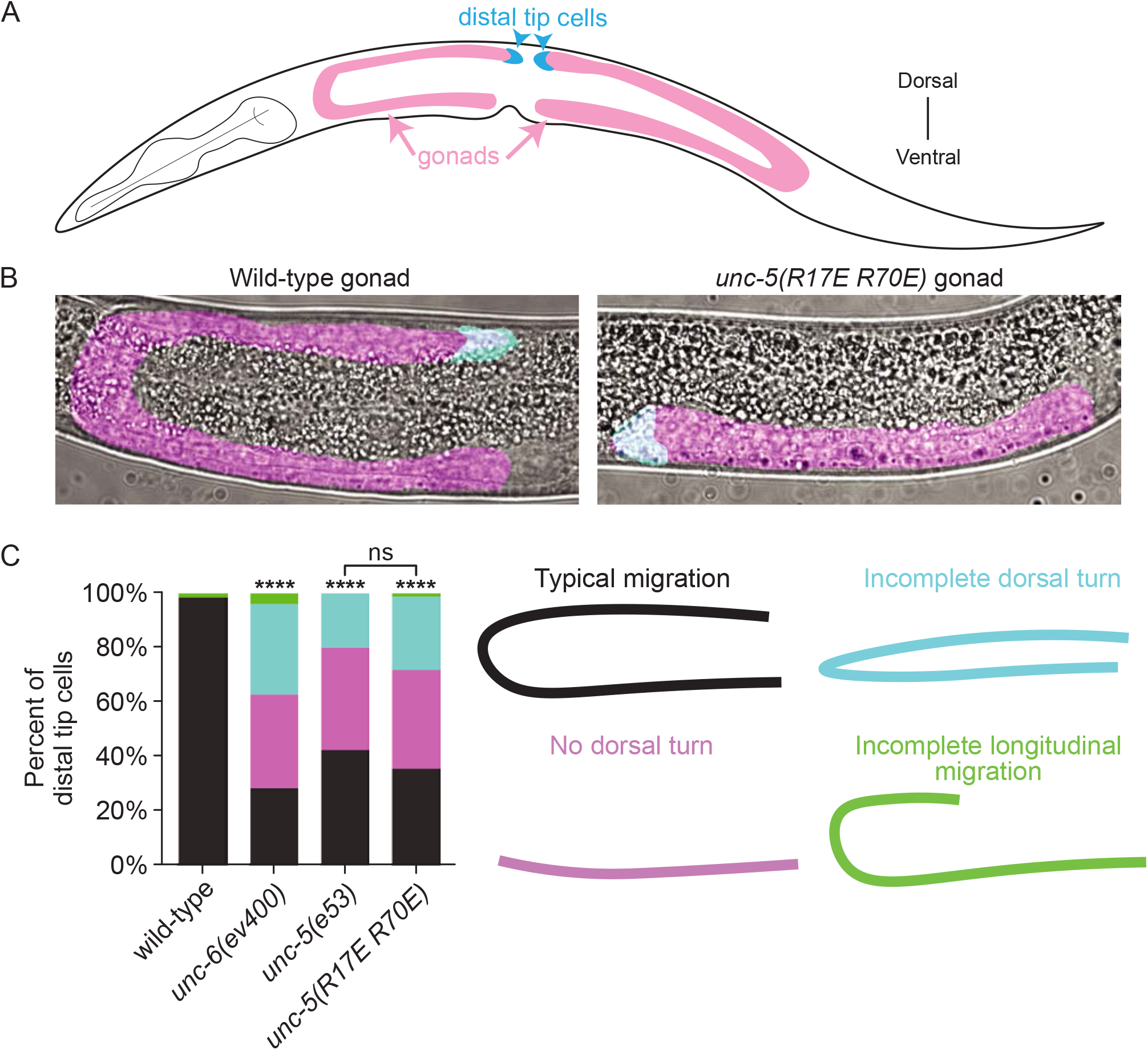
Heparin binding is required for UNC-5-mediated cell migration. **A.** Cartoon of the *C. elegans* gonad. Gonad morphology (magenta) is established by the migration of thedistal tip cell (blue). The ventral-to-dorsal turn of the distal tip cell is dependent on repulsive UNC-5 signaling. **B.** DIC images of gonad morphology in wild-type and *unc-5(R17E R70E)* animals. Blue shaded area denotes the distal tip cell, and the magenta shaded area denotes gonad morphology. **C.** Quantification of distal tip cell migration phenotypes in wild-type, *unc-6(ev400)*, *unc-5(e53)*, and *unc-5(R17E R70E)* animals. Both *ev400* and *e53* are null alleles. Various phenotypes are observed: typical distal tip cell migration (black), a failed dorsal turn (magenta), a partial dorsal turn (blue), and incomplete longitudinal migration (green).

## DISCUSSION

The molecular environment of UNC-5-expressing neurites and cells controls their growth and migrations. In this study, we showed that glycosaminoglycans regulate Netrin–UNC-5 interactions and the processes controlled by them. We deciphered the set of interactions needed to create the Netrin–heparin–UNC-5–UNC-40 complex and show that disrupting interactions between UNC-5 and glycosaminoglycans causes morphological defects during development.

First, we demonstrated experimentally that UNC-5 is a heparin-binding protein. Our structural and biophysical results implicate the boundary of the IG1 and IG2 domains of UNC-5 in its interactions with HS, and possibly with UNC-6, as UNC-5 mutations that break HS and HS-mediated UNC-6 binding also cause small but consistent loss of binding to UNC-6 in the absence of heparin. These findings are supported by previous results showing that the IG domains are necessary and sufficient for binding between human Netrin-1 and UNC5C (Geisbrecht et al., 2003; Kruger et al., 2004) and mouse Netrin-1 and UNC5H2 (Grandin et al., 2016).

We present a model of an UNC-6–heparin–UNC-5 complex that is very large yet well-ordered. For this, we demonstrated that the EGF2 domain of UNC-6/Netrin is a heparin-binding site, and heparin can oligomerize UNC-6 through its secondary interaction site (EGF2 domain), even in the absence of its C-terminal NTR domain, which was identified as the primary heparin interaction site previously (Kappler et al., 2000). Our findings on the EGF2 domain are compatible with previous biochemical studies, which had implicated the Netrin EGF2 domain in Netrin–UNC-5 interactions (Grandin et al., 2016; Kruger et al., 2004), and EGF2 as a potential HS-binding site was recognized (Finci et al., 2014) but had not been shown.

It is reasonable to assume that the length of the linear heparin/HS chains help determine the size of the MDa-sized UNC-6 oligomers and the UNC-6–UNC-5 complex, but our small-angle x-ray scattering data also strongly point to a globular and rigid complex. Considering the flexibility of linear heparin chains, and the still present, but weak, affinity between UNC-6 and UNC-5 in the absence of heparin, we propose that such a rigid and large complex must form via a combination of protein-protein and protein-glycan chains. This composite model of interactions to form Netrin– UNC-5 complexes is appealing, as the specificity of binding may best be provided by protein-protein (UNC-6–UNC-5) contacts, even when interactions with heparin/HS provide much of the binding energy for the formation of the UNC-6–UNC-5 complex.

Since heparin and HS molecules are long, linear chains of sulfated glycans, one such molecule can usually bind more than one protein ligand. However, as we are adding a molar equivalent or excess heparin to UNC-6 or UNC-5+UNC-6 in our experiments, we had expected a majority of UNC-6–heparin or UNC-6–UNC-5–heparin complexes to include one or two protein chains bound to one heparin molecule, rather than very large complexes in the MDa size range. Our observations, therefore, imply that UNC-6 alone or the UNC-6–UNC-5 complex can oligomerize on heparin chains in a cooperative manner, which further supports the model that protein-protein contacts within UNC-6–UNC-6 and UNC-6–UNC-5 oligomers are crucial to support large oligomer formation, in addition to necessary protein contacts with heparin.

While we favor a model where UNC-6–UNC-5 contacts provide specificity for turning on UNC-5-mediated repulsive signaling, it is possible that sulfation domains within the HS chain, as well as additional interactions with the polypeptide chain of an HSPG, can regulate UNC-5 actions and contribute to specificity *in vivo*. This last possibility may be true for an attractive Netrin receptor, *C. elegans* UNC-40/DCC, which was shown to interact with the polypeptide chain of the HSPG LON-2/glypican (Blanchette et al., 2015).

We also attempted to reconstitute the UNC-6–UNC-5–UNC-40 complex, which had been observed before in pull-down experiments (Hong et al., 1999), but not successfully at scale. We saw that heparin was required for a stable complex, probably so that the UNC-6–UNC-5 subcomplex could first form. However, even in the presence of heparin, UNC-40/DCC was only present at a largely substoichiometric ratio, which is unexpected since Netrin–DCC complexes are known to form at a 1:1 ratio, which we have confirmed. This observation supports the hypothesis that UNC-5 and/or heparin might at least partly compete for UNC-40/DCC binding on UNC-6/Netrin (Meijers et al., 2020), which has functional consequences: Repulsive signaling may require UNC-40/DCC to not strongly cluster, which would otherwise lead to attractive signaling, but still be present in a complex with clustered UNC-5 molecules for a repulsive outcome.

In addition to structural insights, we were able to use the molecular reagents, i.e. UNC-5 variants, we engineered to directly ask the following functional question: Is the UNC-5–heparin interaction we discovered necessary for the *in vivo* functions UNC-5 performs, especially those that are UNC-6-dependent? We created animals with the double mutation UNC-5 R17E R70E, which has lost its heparin affinity by 26-fold, and UNC-6ΔC affinity by 62-fold and 2.4-fold in the presence and absence of heparin, respectively, compared to wild-type UNC-5. These animals displayed gonadal morphology defects comparable to *unc-5* and *unc-6* null animals (Hedgecock et al., 1990), supporting our *in vitro* results and structural data on identifying the UNC-5–heparin interaction site, and the functional importance of this interaction.

Our biochemical and structural findings are also in line with the body of available functional data, beyond our own *in vivo* results. It was observed that milder phenotypes of partial *unc-*5 LOF mutants are sensitized in the background of glypican and syndecan mutations, and the repulsive guidance and cell migration actions of UNC-5 depend on LON-2/glypican, functionally linking HSPGs with UNC-5 function (Blanchette et al., 2015; Gysi et al., 2013). The UNC-6 EGF2 domain was identified as essential to UNC-5-mediated functions, as the deletion of the EGF2 domain results in complete loss of repulsive guidance and migration functions (Hedgecock et al., 1990; Lim and Wadsworth, 2002), and R→A mutations on the EGF2 domain causes Netrin-1 to lose its UNC5-mediated anti-apoptopic effect (Grandin et al., 2016). Finally, mutants designed to misfold the IG1 domain of UNC-5 show distal tip cell migration phenotypes and axon guidance effects, although some of the other domain disruptions had similar effects (Killeen et al., 2002). Our work here establishes a model that provides molecular explanations for a large section of the Netrin literature, while enabling future studies using engineered molecules and structural models we have generated, and may serve as a stepping stone towards an atomic-resolution structure of the complete repulsive signaling complex.

## ACKNOWLEDGMENTS

We would like to thank Alexander Jaworski and Michael E. Birnbaum for useful discussions and comments, Hyun Lee and the UIC Biophysics Core Facility for excellent service and support, and Jesse Hopkins and the Biophysics Collaborative Access Team (BioCAT) at the Argonne National Laboratory (ANL) for help with SAXS data collection and processing. This work was supported by the National Institutes of Health grants R01 NS097161 to E.Ö., T32GM138826 to J.M.P., and 1R35GM147179-01 to J.L.M., by the National Science Foundation Graduate Research Fellowship DGE-1656518 to E.L.N., and by the Howard Hughes Medical Institute (HHMI), of which K.S. is an investigator. This research used resources of the Advanced Photon Source, a U.S. Department of Energy (DOE) Office of Science User Facility operated for the DOE Office of Science by ANL under Contract No. DE-AC02-06CH11357. Work at BioCAT was supported by grant P30 GM138395 from the National Institute of General Medical Sciences (NIGMS) of the National Institutes of Health (NIH). This study used resources of the GM/CA@APS, which is supported by the National Cancer Institute (ACB-12002) and the National Institute of General Medical Sciences (NIGMS) (AGM-12006, P30GM138396). This work is also based upon research conducted at the Northeastern Collaborative Access Team beamlines, which are funded by the NIGMS from the NIH (P30 GM124165). The Eiger 16M detector on the 24-ID-E beam line is funded by an NIH-ORIP HEI grant (S10OD021527). This study also used resources of the Berkeley Center for Structural Biology, supported in part by the HHMI, at the Advanced Light Source, a DOE Office of Science User Facility under Contract No. DE-AC02-05CH11231.

## Author Contributions

J.M.P. and J.L.M. designed and executed the directed evolution experiments; J.M.P. and E.Ö. designed and performed structural biology experiments; J.M.P. managed all aspects of protein biochemistry; E.L.N. and K.S. designed and performed in vivo experiments; J.M.P., E.L.N. and E.Ö. prepared the figures, and wrote the manuscript with input from all authors. E.Ö., J.L.M. and K.S. conceptualized the study, and E.Ö. supervised the various aspects of the work.

## MATERIALS AND METHODS

### Protein Expression and Purification

All proteins were expressed using the baculoviral expression system and the lepidopteran High Five cell line in Insect-XPRESS (Lonza, cat. no. BELN12-730Q). Baculoviruses were produced using homologous recombination in insect cells by co-transfection with linearized baculoviral DNA and the transfer vector pAcGP67A (BD Biosciences), which carries an N-terminal gp64 signal peptide for secretion. All proteins were tagged C-terminally with a hexahistidine tag for purification. When indicated, some constructs were tagged C-terminally with a BirA recognition sequence (GLNDIFEAQKIEWHE), for facile enzymatic biotinylation, followed by a hexahistidine tag. Biotinylation was performed using the BirA ligase reaction kit (Avidity, cat. no. BirA500).

UNC-5, UNC-6 and UNC-40 constructs of various lengths were purified first with Ni-NTA metal-affinity chromatography (QIAGEN, cat. no. 30210) from conditioned media, followed by size-exclusion chromatography with either Superose 6 Increase 10/300 (GE Healthcare, cat. no. 29-0915-96) or Superdex 200 Increase 10/300 columns (GE Healthcare, cat. no. 28-9909-44). UNC-5 and UNC-40 are purified in 10 mM HEPES pH 7.2 and 150 mM NaCl (HEPES-buffered saline, HBS), while UNC-6ΔC exhibits aggregation and stickiness to chromatography resin at physiological pH, as recently recognized by another group (Krahn et al., 2019). We tested numerous buffer conditions in size-exclusion chromatography experiments, and observed that UNC-6ΔC can be solubilized in a buffer with 500 mM NaCl, and shows the best elution profile and protein recovery when in 10 mM HEPES pH 7.2, 150 mM NaCl, 100 mM MgSO_4_ (HBS-MS), presumably as magnesium acting as a chaotrope and sulfate interacting with the heparin-binding site(s) (Figure S1B).

### Yeast Surface Display

All yeast clones and libraries were made using electrically competent EBY100 yeast (Thermo Fisher, cat. no. 83900) (Chao et al., 2006). UNC-5 ECD, containing 2 Ig and 2 Tsp1 domains, was displayed on yeast via fusion of the C-terminus of UNC-5 ECD to the N-terminus of Aga2. A Myc-tag was included on the C-terminus of UNC-5 for quantification of fusion protein expression via staining with an Alexa Fluor 488-coupled anti-myc (Thermo Fisher, cat. no. MA1-980-A488). To test binding to UNC-6 of yeast displaying UNC-5, an UNC-6 construct lacking the C-terminal NTR domain, known to cause aggregation, and containing a BirA biotin ligase recognition sequence was expressed and purified (UNC-6ΔC). A yeast staining reagent was created by biotinylating UNC-6ΔC and incubating it with Alexa Fluor 647 (Thermo Fisher, cat. no. A20347)-coupled streptavidin.

The UNC-5 ECD display library was created using random error-prone mutagenesis. Separation of yeast populations was done via magnetic-activated cell sorting (Miltenyi Biotec, cat. no. 130-091-051) using either streptavidin (Miltenyi Biotec, cat. no. 130-048-101) or anti-Alexa Fluor 647 microbeads (Miltenyi Biotec, cat. no. 130-091-395). Progress during selections was assessed by UNC-6ΔC binding and monitored using flow cytometry on a BD Accuri C6 flow cytometer.

The yeast library underwent three rounds of selection, performed in decreasing concentrations of heparin from porcine intestinal mucosa (Sigma-Aldrich, cat. no. 9041-08-1). The first round was performed with streptavidin microbeads in 400 nM UNC-6ΔC monomers and 4 μg/ml heparin. The second round was performed with anti-Alexa Fluor 647 microbeads in 125 nM UNC-6ΔC monomers and 0.062 μg/ml heparin. The third round was performed the same as the second round, excluding heparin.

Individual yeast clones (96 total) were isolated after the third selection round. These clones were tested for ability to bind UNC-6ΔC. Of the 96 clones, 8 higher affinity clones were chosen for further analysis. Binding affinities to UNC-6ΔC on yeast without added heparin were measured and the DNA sequence of UNC-5 ECD was sequenced for each clone.

### Protein Crystallization

A construct containing the first two IG domains of *C. elegans* UNC-5 with a C-terminal hexahistidine tag was purified as described above and concentrated to 25 mg/ml in HBS. Crystals were grown with the sitting-drop vapor diffusion method, and using microseeding to improve single crystal growth in the crystallant 1.25 M (NH_4_)_2_SO_4_, 0.1 M Tris pH 8.5, 0.2 M Li_2_SO_4_ at 22°C. Cryoprotection was achieved using a cryoprotectant supplemented with 30% Glycerol by flash-freezing in liquid nitrogen.

UNC-5 IG1-2 with the short heparin fragment dp4 (Iduron, cat. no. HO04; molecular weight, ~1.2 kDa) was crystallized after mixing purified protein with dp4 at a 1:3 molar ratio. Crystallization was achieved with 20 mg/ml UNC-5 using vapor diffusion with the hanging-drop method with the crystallant 4% (v/v) Pentaerythritol ethoxylate (3/4 EO/OH), 0.1 M sodium acetate pH 4.6, 16% (w/v) PEG 8000, 0.2 M NaCl at 22°C. Crystals were cryoprotected in 0.1 M sodium acetate pH 4.8, 16% PEG 8000, 0.2 M NaCl, 25% ethylene glycol, 60 μg/ml dp4 by flash-freezing in liquid nitrogen.

UNC-6ΔC with a C-terminal hexahistidine tag was purified over Superdex200 10/300 after being mixed with heparin-dp16 (Iduron, cat. no. HO16; molecular weight, ~4.65 kDa) at a 1:1.5 molar ratio in the HBS-MS buffer. Protein was concentrated to x mg/ml; we could not check if any dp16 was carried over during purification and concentration. Crystallization was achieved using hanging-drop vapor diffusion with 0.1 M Tris, pH 8.0, 10% (w/v) Polyethylene glycol 8,000 at 22°C. Crystals were cryoprotected with 0.1 M Tris pH 8, 10% PEG 8,000, 30% glycerol and 0.12 mg/ml dp16. We did not observe any electron density for any heparin-like molecules, as crystal packing appears to be blocking much of our predicted HS-binding site, except for an ordered putative sulfate ion (present at 100 mM in the protein buffer).

### Structure Determination by X-ray Crystallography

Crystallographic images were indexed and scaled using the *XDS* package programs (Kabsch, 2010). Molecular replacement was performed with *PHASER* (McCoy et al., 2007) in the *PHENIX* package (Liebschner et al., 2019). UNC-5 IG1+2 structure was solved using the structure of the human Unc5A ectodomain (Seiradake et al., 2014) with domains 1 and 2 as separate molecular replacement models (PDB ID: 4V2A) after homology modeling for the *C. elegans* sequences with *MODELLER* (Webb and Sali, 2016). The structure with heparin-dp4 was determined using our UNC-5 IG1+2 structure as the molecular replacement model. UNC-6ΔC structure was solved using the AlphaFold model as the molecular replacement model (Varadi et al., 2022). *phenix.refine* was used to refine the model structure in reciprocal space (Afonine et al., 2012), and *COOT* was used for model building and correction in real space (Emsley et al., 2010). Model building and refinement was guided by *MOLPROBITY* tools within *PHENIX* for checking chemical geometry (Chen et al., 2010).

The diffraction data for the UNC-5 IG1+2 with Heparin-dp4 crystals were highly anisotropic (See Table 1 for details). The dataset was reduced using the XDS package, and then ellipsoidically truncated using the STARANISO server (Tickle et al., 2018), which was used during refinement.

For analysis of sequence and structural features, we heavily used the APBS plugin in PyMOL for electrostatic potential surface calculations (Jurrus et al., 2018), and ConSurf for calculating sequence conservation (Ashkenazy et al., 2016). Isoelectric points and extinction coefficients were calculated using the Protparam server (Gasteiger et al., 2005).

### Surface Plasmon Resonance

A Biacore T200 (GE Healthcare) at 22°C was used to measure the binding affinities by the equilibrium method, as kinetics of binding was too fast to measure in most occasions. Biotinylated UNC-6ΔC or heparin (Millipore Sigma, cat. no. 375054) was captured on a streptavidin-coated sensor chip (Cytiva, cat. no. 29104992). Analytes for biotinylated UNC-6ΔC included UNC-5 ECD (with and without 5 μg/ml heparin) and heparin. Analytes for biotinylated heparin included UNC-5 ECD and UNC-6ΔC. The running buffer for all SPR experiments was 10 mM HEPES pH 7.2, 150 mM NaCl, 0.05% Tween20, 0.1% BSA, supplemented with either 20 or 50 mM MgSO_4_ when UNC-6ΔC was an analyte to reduce nonspecific binding. A regeneration step was performed after each injection using 10 mM HEPES pH 7.2, 150 mM NaCl, 0.05% Tween20, 100 mM MgSO_4_.

Using Biacore evaluation software, the dissociation constant (KD) was calculated by fitting the equilibrium data with a 1:1 binding model. whenever binding was observed to nearly saturate. With mutants that had very low affinities saturation of the sensor surface was impossible; therefore, maximum SPR response (i.e. *R*_max_ at saturation) was restrained to WT Rmax values for *K*_D_ calculations in Prism (GraphPad).

### Small-angle X-ray Scattering

Small-angle x-ray scattering data was collected at the APS beamline 18-ID using a Superose 6 10/300 column on the SEC-SAXS setup. The protein samples were prepared by mixing UNC-6ΔC, UNC-5 ECD and UNC-40 ECD at molar equal ratios with heparin at 1.5-times the molar equivalent. No heparin was present in the running buffers. Detailed experimental conditions and parameters are listed in Table S4. Data was processed in BioXTAS RAW version 2.1.4 (Hopkins et al., 2017), using tools from the ATSAS package, version 3.1.1 (Manalastas-Cantos et al., 2021). For the UNC-6–heparin–UNC-5 data, we used evolving factor analysis (EFA) deconvolution as implemented in RAW to extract scattering from the complex and not the isolated components as a precaution (Meisburger et al., 2016). However, analysis of frames corresponding to just those collected for the major peak gave very similar results to those we report using EFA in Table S4.

### Generation of mutant *C. elegans* with CRISPR/Cas9 and phenotypic analysis of animals

*C. elegans* were grown at 20°C on NGM plates and fed with *Escherichia coli* according to standard procedures (Brenner, 1974). N2 Bristol was used as the wildtype reference strain. The following additional strains were used in this study: *unc-6(ev400)* X (NW434), *unc-5(e53)* IV (TV2348), and *unc-5(R17E R70E)* VI.

Base pair mutations in *unc-5* were generated by microinjections of CRISPR/Cas9 protein complexes in N2 Bristol animals. Genome editing using CRISPR-Cas9 was carried out by standard protocols (Dokshin et al., 2018). Cas9 protein and tracRNA (IDT) were injected at a concentration 1.525μM each. crRNAs (IDT) were injected at a concentration of 1.525 μM. Single-stranded DNA repair templates were ordered as oligomers (ThermoFisher) and injected at μM. pRF6(*rol-6(su1006)*) was used as a co-injection marker and selected against after confirmation of a successful genome edit. Confirmation of genome edits was performed by sequencing of F2 animals. The R17E and R70E edits were generated sequentially. The R17E mutation was generated with the following crRNA sequence: CTTATTTTCAATGACATAAC. The R70E mutation was generated with the following crRNA sequence: GAATCGACGTAGACACATC.

*C. elegans* hermaphrodites in the late L4 and young adult stages were screened for gonad morphology and distal tip cell migration. Animals were anesthetized using 6mM levamisole (Sigma-Aldrich) in M9 buffer and mounted on 4% agarose pads. DIC images were captured using an inverted Zeiss Axio Observer Z1 microscope equipped with a Prime 95B Scientific CMOS camera (Photometrics). 3i Slidebook software was used to control imaging. A water-immersion C-Apochromat 40x 0.9 NA objective was used for all imaging. All images were processed using ImageJ/Fiji (National Institutes of Health). Pseudocoloring of gonad morphology and distal tip cells were performed manually using the freehand selector tool. The resulting regions of interest were used to generate masks which were merged with the original DIC image.

## SUPPLEMENTAL FIGURE LEGENDS

**Figure S1. (related to Figure 1.).**
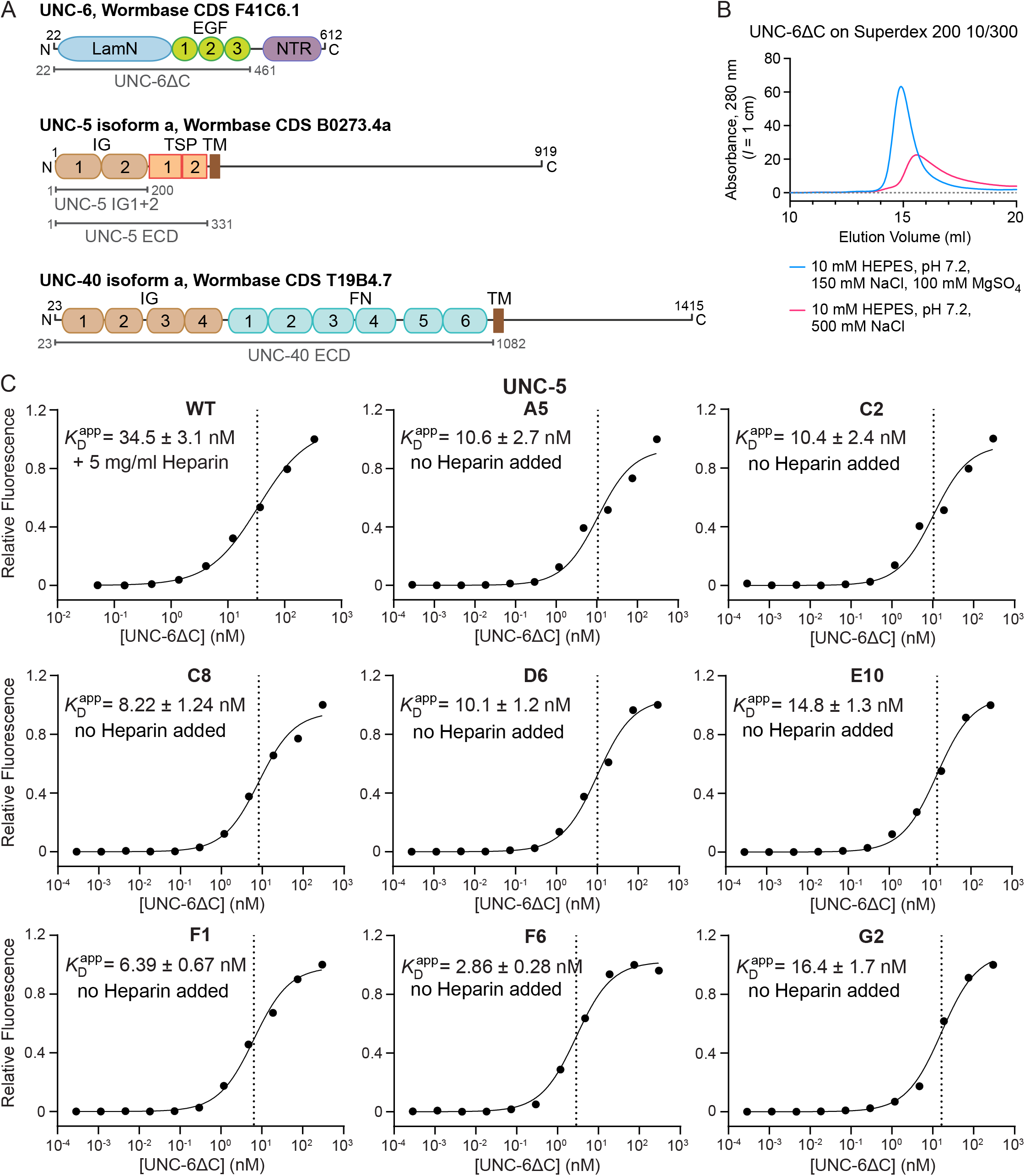
**A.** Schematic for boundaries of constructs used for binding studies, crystallography, biophysics, and directed evolution. **B.** Size-exclusion chromatograms of hexahistidine-tagged UNC-6ΔC run in HBS (red), HBS containing 500 mM NaCl (blue), and HBS-MS (green). The column is a Superdex 200 10/300 (GE Healthcare). 100 μg was loaded in both runs: The run with HBS-MS buffer recovered 89.8 μg as measured by the area under the peak (76.11 mAU*ml), while the run with high-NaCl HBS showed stronger signs of non-specific stickiness to the resin, allowing for 66.4 μg to be recovered (area = 56.25 mAU*ml). Abs-0.1% value (absorbance at 1 mg/ml) used for UNC-6ΔC is 0.845.**C.** Binding isotherm for UNC-6ΔC to yeast expressing WT UNC-5 ECD in the presence of 5 μg/ml heparin and to eight yeast colonies expressing UNC-5 ECD variants without added heparin.

**Figure S2 (related to Figure 2.).**
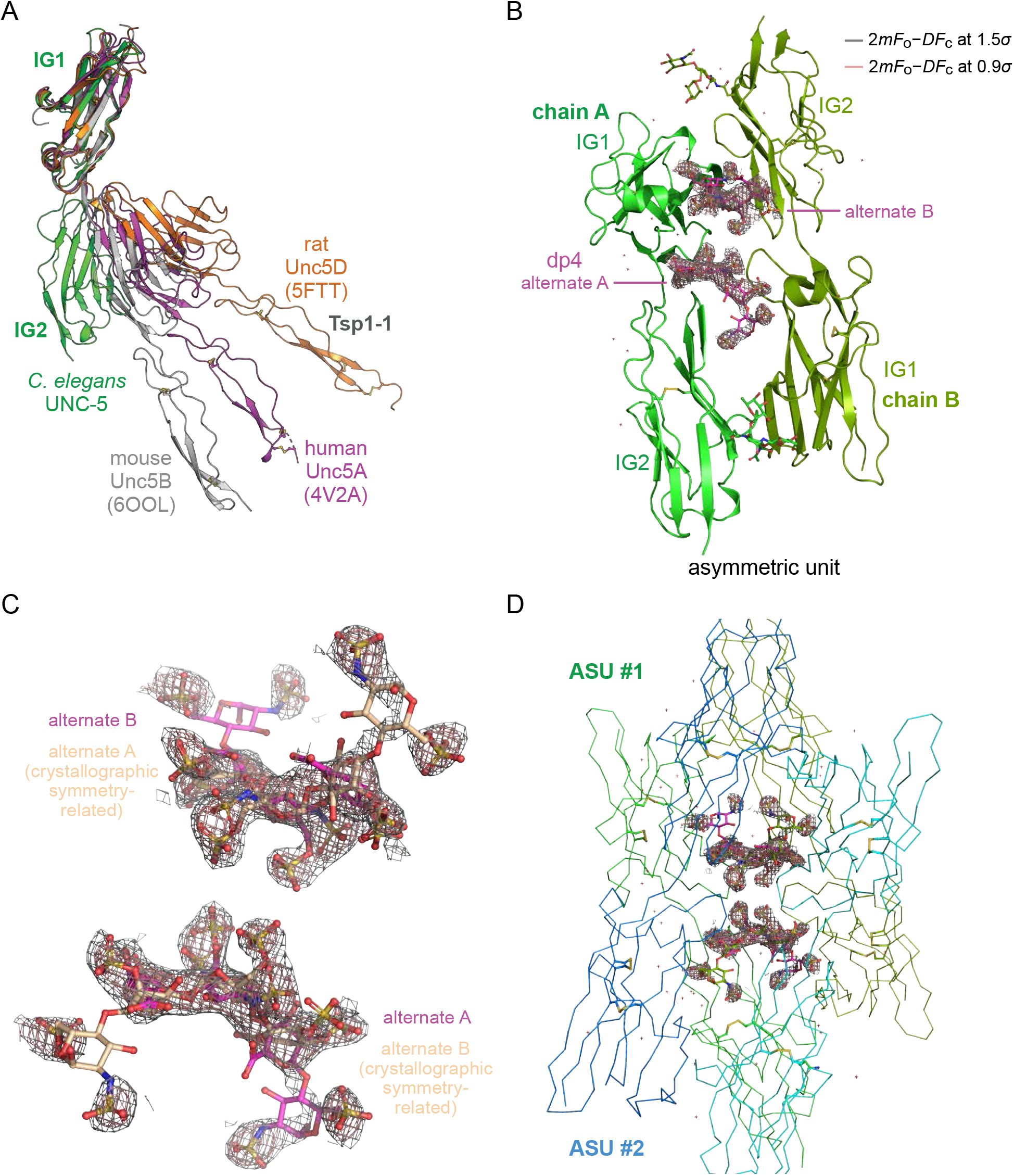
**A.** Superposition of reported UNC-5 structures in the literature and our UNC-5 IG1-2 structure, where the IG1 domains are overlayed, demonstrating wide flexibility of the IG1-IG2 boundary. **B.** The asymmetric unit of the UNC-5-dp4 crystals, with 2*mF*o−*DF*c density carved around the dp4 ligand. Heparin-dp4 was modeled as two alternate conformations in equivalent positions of the two UNC-5 chains related by non-crystallographic symmetry (light and dark green). **C.** Electron density for heparin-dp4 with dp4 from two neighboring asymmetric units shown (colored magenta and salmon), each with two alternate copies. Density is strongest at the electron-rich sulfates. **D.** Electron density for dp4 surrounded by four UNC-5 chains, contributed by two asymmetric units.

**Figure S3 (related to Figure 3.).**
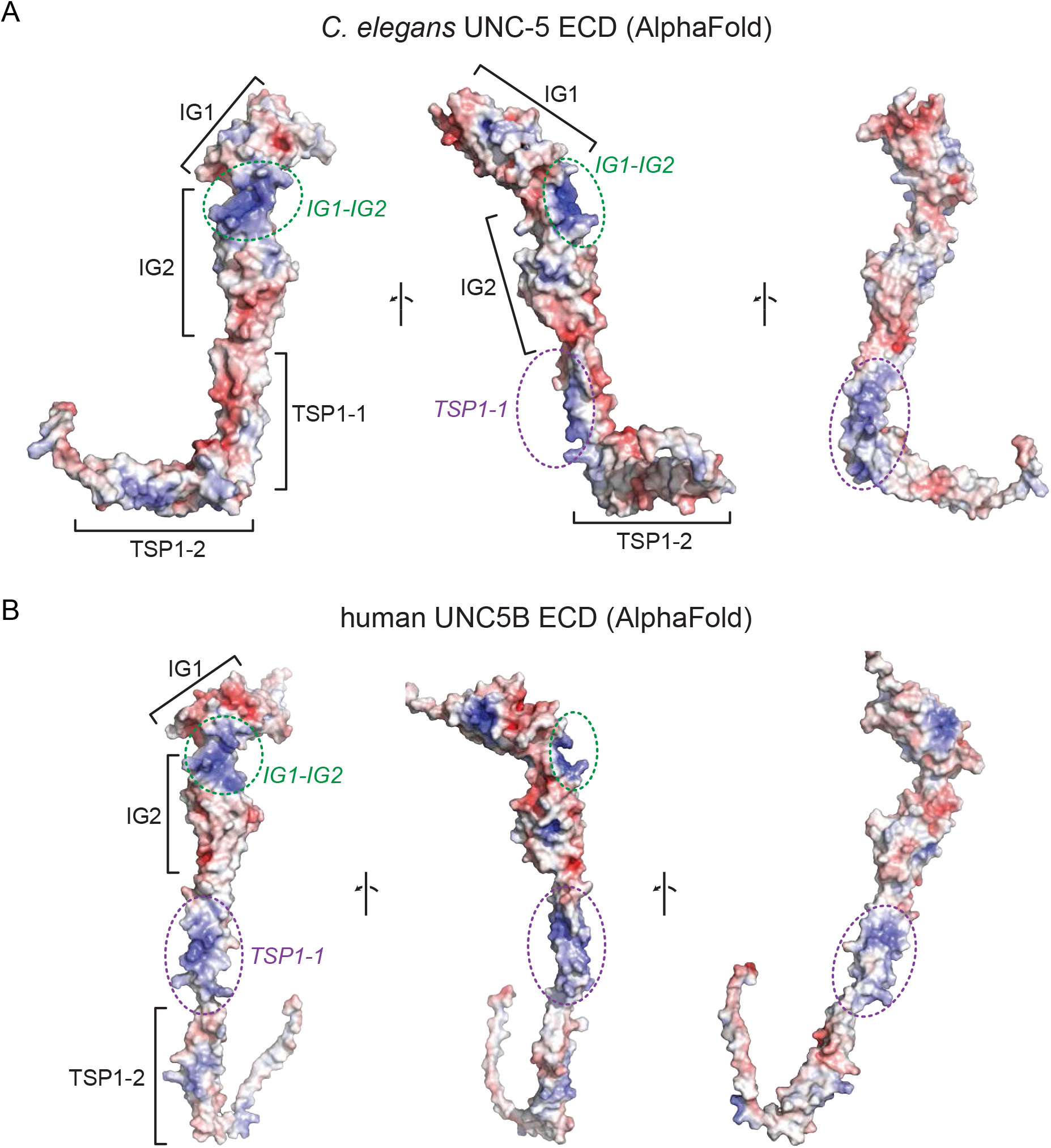
Electrostatic potential surfaces for *C. elegans* UNC-5 **(A)**, and human UNC5B **(B)**. Structural models for the entire ectodomains were taken from EBI’s AlphaFold database. Both show positively charged surfaces on the IG1-IG2 boundary and on the first Tsp1 domain; however, the IG1-IG2 boundary is more conserved (Figure 3B).

**Figure S4. (related to Figure 4.).**
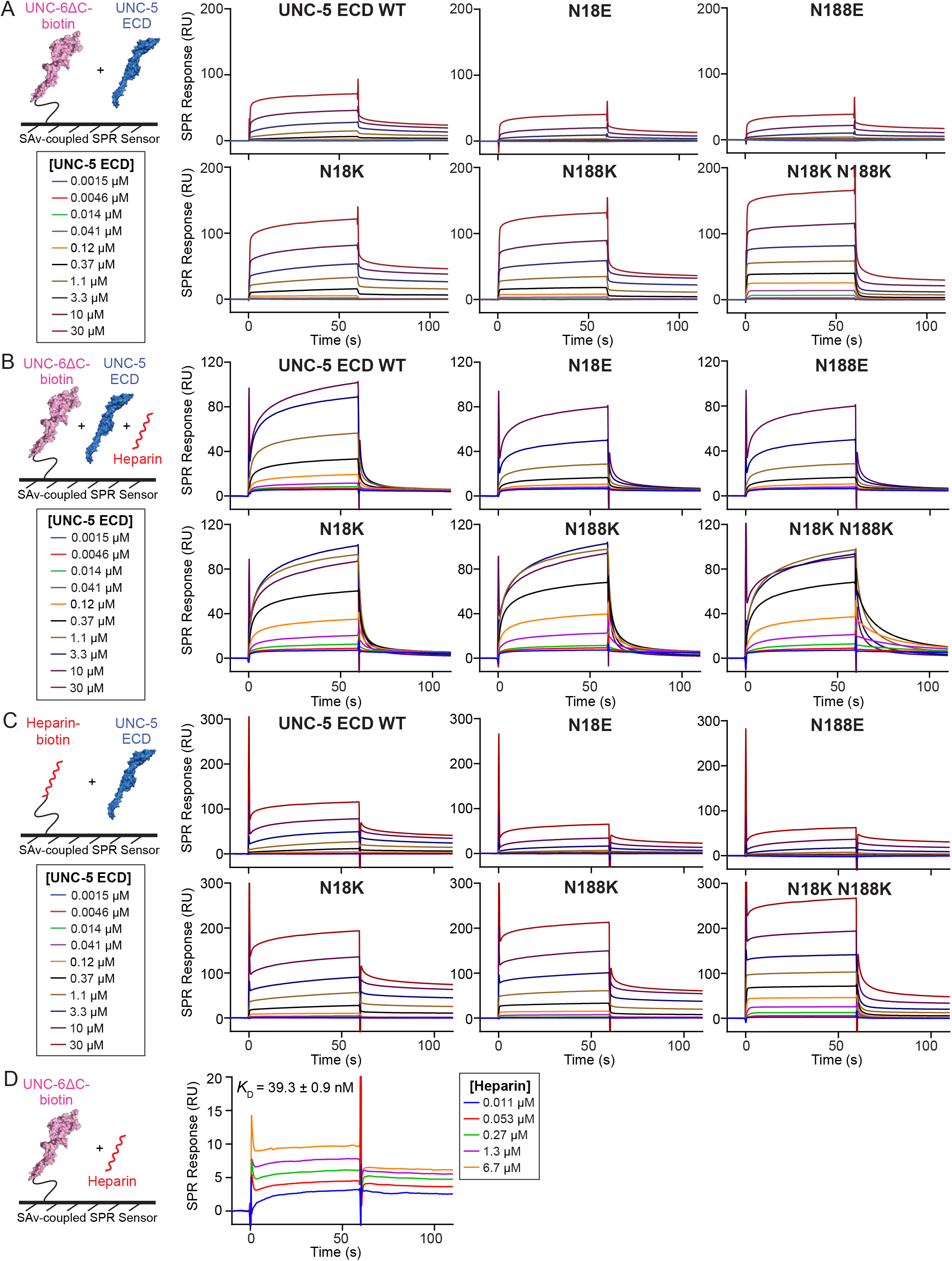
SPR sensorgrams for isotherms reported in Figure 4. Same channel on the SPR chip was used within each panel, allowing for direct comparison of response values between mutants.

**Figure S5. (related to Figure 5.).**
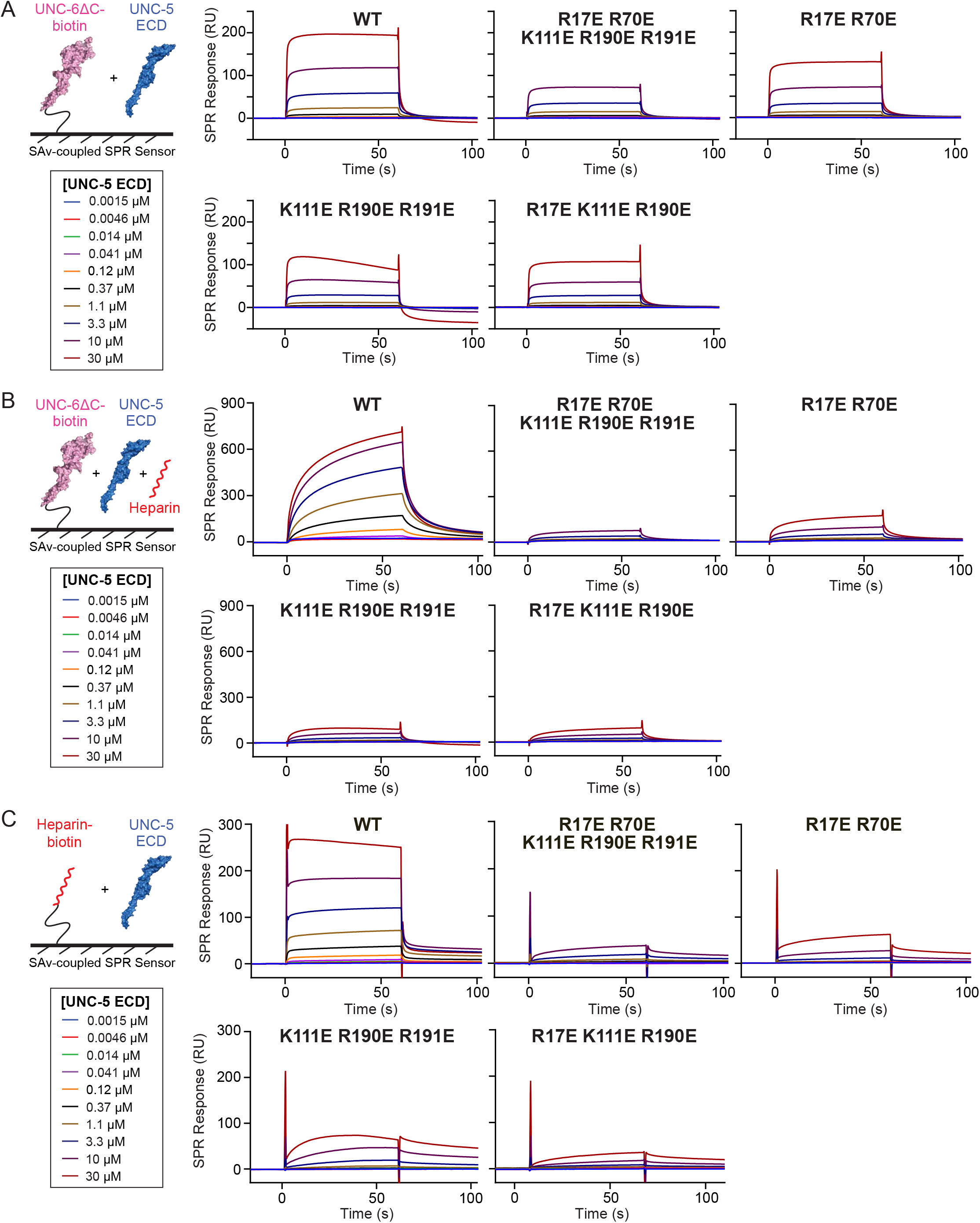
SPR sensorgrams for isotherms reported in Figure 5. Same channel on the SPR chip was used within each panel, allowing for direct comparison of response values between mutants.

**Figure S6 (related to Fig. 6.).**
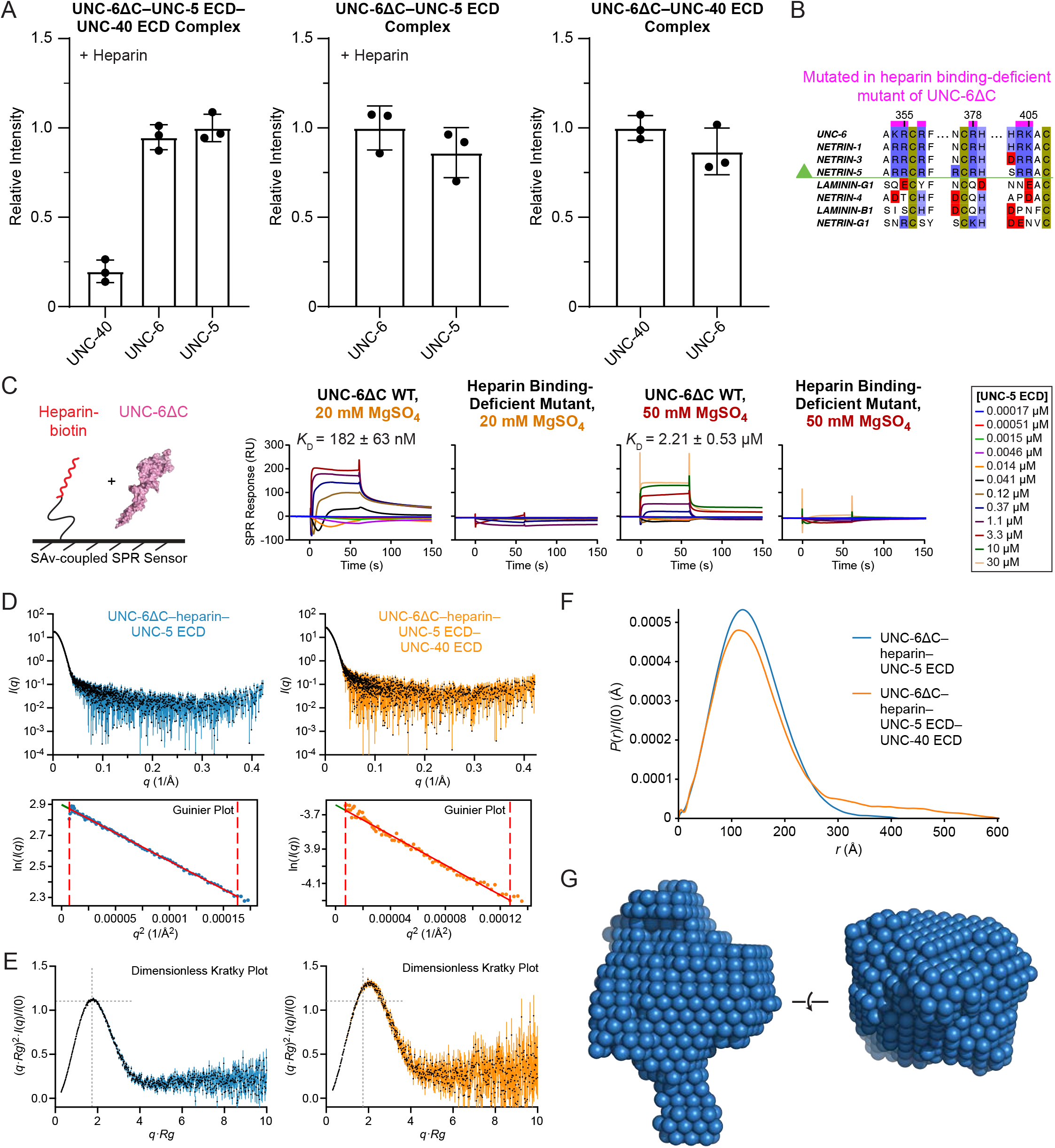
**A.** Coomassie blue staining intensities of purified UNC-6ΔC–UNC-5 ECD–UNC-40 ECD (left), UNC-6ΔC–UNC-5 ECD (middle), and UNC-6ΔC–UNC-40 ECD (right) complexes were analyzed (n = 3 lanes in SDS-polyacrylamide gels). All complexes were purified in SEC runs after mixing proteins in equimolar ratios, and with excess heparin (1.5 molar ratio) for the first two complexes. Band intensities were quantified using Image Lab (Bio-Rad Laboratories) and analyzed in Prism (GraphPad). **B.** Partial sequence alignment of UNC-6 and human Netrin homologs. **C.** SPR sensorgrams for isotherms reported in Figure 6. **D.** Scattering intensity and Guinier plots for SEC-SAXS data collected for complex peaks for the UNC-6–heparin–UNC-5 (blue) and UNC-6–heparin–UNC-5–UNC-40 samples (orange). **E.** Dimensionless Kratky plots for SAXS data. **F.** Pair distribution plots for SAXS data, showing a globular shape for the UNC-6–heparin–UNC-5 (blue) complex and the UNC-6–heparin–UNC-5–UNC-40 complex (orange). **G.** Bead model calculated by DAMMIF’s *damfilt* function (ATSAS package) for the UNC-6–heparin–UNC-5 complex.

**Table S1 (related to Figures 2 and 6.)** Data and refinement statistics for x-ray crystallography.

**Table S2 (related to Figure 4).**
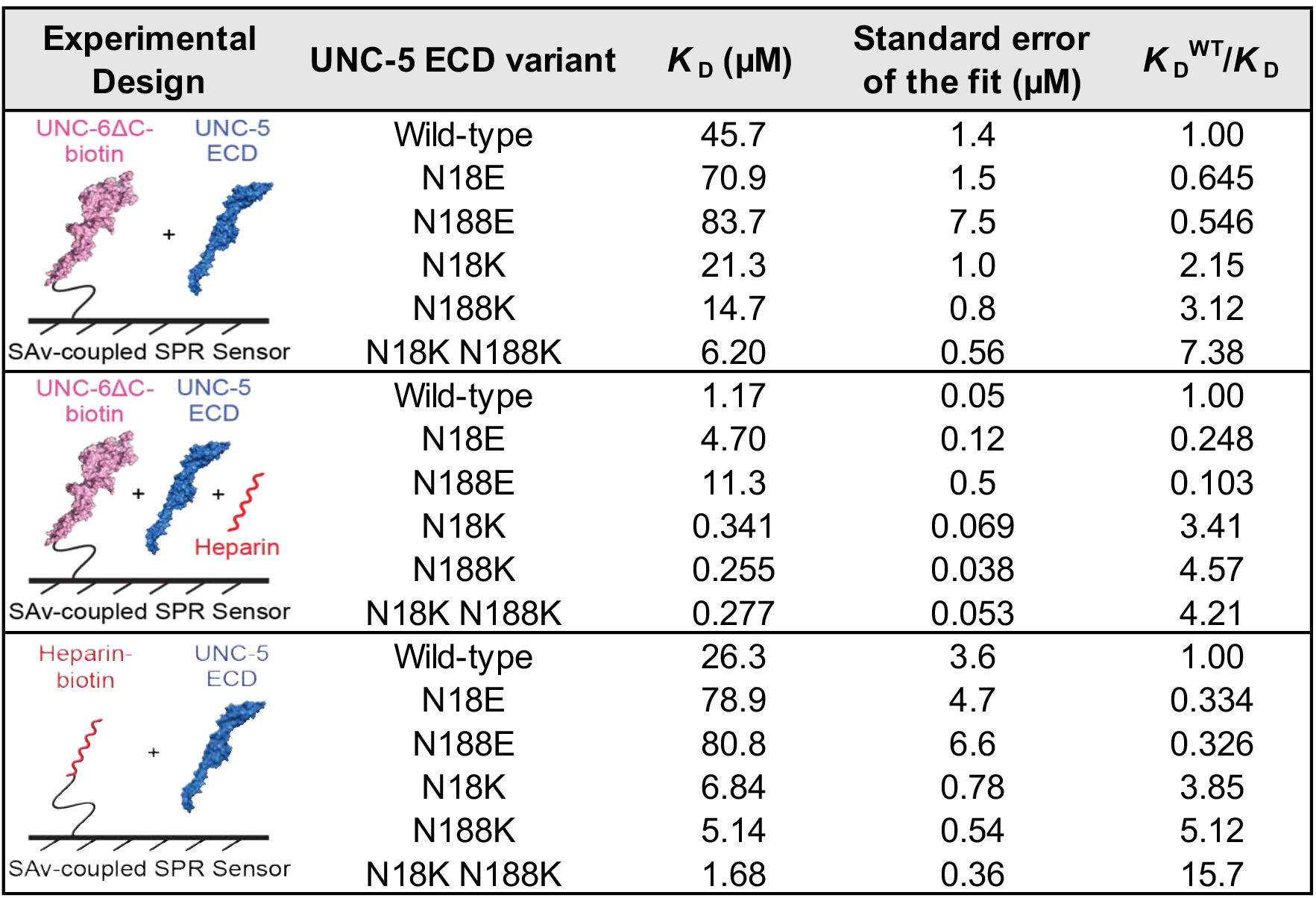
Summary of fitting of SPR sensorgrams in Figure 4, with standard error of the parameter reported by Prism (GraphPad).

| Experimental Design | UNC-5 ECD variant | $K_D$ (μM) | Standard error of the fit (μM) | $K_D^{WT}/K_D$ |
| --- | --- | --- | --- | --- |
| SAv-coupled SPR Sensor | Wild-type | 45.7 | 1.4 | 1.00 |
|  | N18E | 70.9 | 1.5 | 0.645 |
|  | N188E | 83.7 | 7.5 | 0.546 |
|  | N18K | 21.3 | 1.0 | 2.15 |
|  | N188K | 14.7 | 0.8 | 3.12 |
|  | N18K N188K | 6.20 | 0.56 | 7.38 |
| SAv-coupled SPR Sensor | Wild-type | 1.17 | 0.05 | 1.00 |
|  | N18E | 4.70 | 0.12 | 0.248 |
|  | N188E | 11.3 | 0.5 | 0.103 |
|  | N18K | 0.341 | 0.069 | 3.41 |
|  | N188K | 0.255 | 0.038 | 4.57 |
|  | N18K N188K | 0.277 | 0.053 | 4.21 |
| SAv-coupled SPR Sensor | Wild-type | 26.3 | 3.6 | 1.00 |
|  | N18E | 78.9 | 4.7 | 0.334 |
|  | N188E | 80.8 | 6.6 | 0.326 |
|  | N18K | 6.84 | 0.78 | 3.85 |
|  | N188K | 5.14 | 0.54 | 5.12 |
|  | N18K N188K | 1.68 | 0.36 | 15.7 |

**Table S3 (related to Figure 5).** Summary of fitting of SPR sensorgrams in Figure 5, with standard error of the parameter reported by Prism (GraphPad).

| Experimental Design | UNC-5 ECD variant | $K_D$ (μM) | Standard error of the fit (μM) | $K_D^{WT}/K_D$ |
| --- | --- | --- | --- | --- |
| SAV-coupled SPR Sensor | Wild-type | 3.86 | 0.34 | 1.00 |
|  | R17E R70E K111E R190E R191E | 62.3 | 3.3 | 0.0620 |
|  | R17E R70E | 100 | 3 | 0.0385 |
|  | K111E R190E R191E | 77.0 | 8.8 | 0.0502 |
|  | R17E K111E R190E | 197 | 11 | 0.0196 |
| SAV-coupled SPR Sensor | Wild-type | 12.8 | 0.2 | 1.00 |
|  | R17E R70E K111E R190E R191E | 27.1 | 0.9 | 0.473 |
|  | R17E R70E | 30.8 | 1.1 | 0.416 |
|  | K111E R190E R191E | 52.5 | 5.1 | 0.244 |
|  | R17E K111E R190E | 43.0 | 2.1 | 0.298 |
| SAV-coupled SPR Sensor | Wild-type | 1.44 | 0.11 | 1.00 |
|  | R17E R70E K111E R190E R191E | 77.6 | 10.9 | 0.0186 |
|  | R17E R70E | 89.6 | 8.1 | 0.0161 |
|  | K111E R190E R191E | 178 | 25 | 0.00809 |
|  | R17E K111E R190E | 187 | 17 | 0.00768 |

**Table S4 (related to Figure 6.)** Data statistics and analysis for small angle x-ray scattering experiments.

